# Intra-thalamic coordination maintains arousability in slow-wave sleep

**DOI:** 10.64898/2026.11.15.766872

**Authors:** Junrong Han, Qiuyou Xie, Ying Huang, Man Li, Xiaoqing Hu, Georg Northoff, Yuanqing Li, Jun Zhang, Pengmin Qin

## Abstract

Slow-wave sleep preserves the capacity for spontaneous or stimulus-evoked awakening despite reduced responsiveness, yet how thalamic nuclei coordinate to preserve arousability remains unclear. We analysed six fMRI and EEG datasets spanning sleep, propofol anesthesia and unresponsive wakefulness syndrome (UWS). During N2 and N3, centromedian (CM) and ventral posterolateral (VPL) nuclei showed convergent edge-community profiles and increased closed-loop intra-thalamic motifs, changes not observed during anesthesia or UWS. During N3, directed effective connectivity along CM→VPL→pulvinar (PUL) increased versus wakefulness and N2. Stronger CM→VPL and VPL→PUL connectivity covaried with greater N3 delta power, which was higher before awakening than before transitions to N2 and positively associated with three-month improvement in behavioural responsiveness in UWS. These findings identify a CM–VPL–PUL-centred pattern of intra-thalamic organization linked to preserved arousability during slow-wave sleep, extending accounts focused on individual nuclei and thalamocortical interactions.

## Introduction

Within the distributed cortico-subcortical system that supports consciousness, the thalamus has been increasingly identified as a key hub^1–6^, yet how its diverse nuclei interact to sustain consciousness remains a central challenge in neuroscience. Three conditions are widely used to probe the neural substrates of consciousness: slow-wave sleep (also termed N3 sleep), general anesthesia, and unresponsive wakefulness syndrome (UWS)^7–22^. Among these states, slow-wave sleep is distinctive in preserving arousability: despite markedly reduced responsiveness to external stimuli, individuals retain the capacity for spontaneous or stimulus-evoked awakening^23^. By contrast, this capacity is markedly impaired during general anesthesia and in UWS^24–27^. Anesthesia reflects a pharmacologically induced loss of consciousness ^28–30^, whereas UWS is a pathological condition in which patients exhibit wake-like behaviours (e.g., eye opening) without awareness (formerly known as the vegetative state) ^26,27,31^. However, the neural mechanisms underlying preserved arousability during slow-wave sleep remain unclear.. Recent evidence indicates that thalamic nuclei show temporally ordered activity changes preceding spontaneous awakening, suggesting that coordinated interaction among nuclei may help maintain arousability and prepare the brain for awakening during sleep ^21^.

Prior research on sleep–wake regulation has emphasized distributed cortical and subcortical systems, including thalamocortical interactions and arousal-related neuronal populations in the thalamus, hypothalamus, basal forebrain and brainstem^32–35^. Within this framework, the thalamus has been proposed as a key hub coordinating arousal-related processing between cortical and subcortical systems^36–38^, and specific thalamic nuclei have been implicated in sleep–wake regulation^5,39,40^. However, to date most studies have focused on thalamocortical pathways or single nuclei, leaving unclear how interactions among distinct thalamic nuclei would preserve arousability during slow-wave sleep.

Stimulatin<u>g</u> intralaminar nuclei, such as centromedian nucleus (CM), has been associated with enhanced cortical engagement and recovery of behavioral responsiveness in asleep/anesthetized animal models^3,10,41–44^ and in disorders of consciousness (DOC), including UWS^45–49^. However, intralaminar thalamic stimulation does not reliably restore full consciousness in patients with DOC^47,48,50–54^, suggesting that broader thalamic coordination and thalamocortical connection may be required. Consistent with this view, nuclei beyond the intralaminar group—such as the ventral posterolateral nucleus (VPL) and the pulvinar (PUL)—have also been linked to changes in global co-activation during deep sleep^5^. Notably, evidence of an orderly temporal activation sequence across thalamic nuclei during spontaneous awakenings from sleep^21^ implies that interaction between nuclei may be a key mechanism supporting sleep arousability. Together, these findings support a subnetwork view of thalamus: multiple nuclei can share transcriptional, morphological, physiological, and connectivity profiles, and jointly form a subnetwork for serving a specific function^55^. From this perspective, consciousness may depend on coordinated interactions rather than the activity of any single nucleus in isolation. Accordingly, a key question is how thalamic nuclei interact to preserve the capacity for awakening (i.e., arousability) in slow-wave sleep, and whether such coordination is disrupted or absent in anesthesia and UWS.

To investigate whether preserved arousability involves coordinated thalamic subnetworks, we sought to characterize two complementary aspects of intra-thalamic organization: overlapping functional architecture and directed interactions. Edge-centric analysis characterizes overlapping community structure by clustering temporally co-fluctuating connections, allowing individual thalamic nuclei to participate in multiple functional communities^56–58^. Complementing this approach, neural perturbational inference (NPI) estimates directed effective connectivity among thalamic nuclei by quantifying responses to virtual perturbations of a neural-network surrogate of brain dynamics^59^.

A further challenge is to determine whether intra-thalamic coordination during sleep is specifically related to arousability, rather than simply reflecting the neurophysiological features of sleep itself. Slow-wave sleep provides a particularly informative stage for addressing this question because sensory responsiveness is markedly reduced while arousability is preserved^23^. Slow-wave sleep is also characterized by prominent slow-wave activity, commonly indexed by delta activity and associated with highly synchronized neuronal dynamics^23^. Although delta activity has been attributed primarily to neocortical mechanisms^60–62^, growing evidence indicates that thalamic nuclei contribute to its generation, synchronization and propagation^44,63–65^. In addition, while delta activity has traditionally been associated with diminished consciousness, accumulating evidence suggests that prominent delta activity can occur in conscious states and transiently increase during arousal or awakening from slow-wave sleep^18,66^. We therefore hypothesized that, if intra-thalamic coordination during slow-wave sleep supports preserved arousability, its strength might covary with delta power, and delta power would then be associated with the propensity for subsequent awakening.

To address above questions, we combined fMRI and EEG across multiple datasets to investigate the thalamic mechanisms linked to preserved arousability in slow-wave sleep (Fig. 1A). We first analysed three fMRI discovery datasets spanning sleep, propofol anesthesia and UWS to identify changes in intra-thalamic organization and directed interactions during sleep that were not observed during deep sedation or in UWS. We then tested the robustness of these findings in an independent sleep fMRI dataset. Next, using simultaneous EEG-fMRI data, we examined whether the identified thalamic directed interactions covaried with N3 delta power. We further assessed the behavioural relevance of N3 delta power by comparing N3 segments followed by wakefulness (N3F-W) with those followed by N2 sleep (N3F-N2), and its clinical relevance by examining the association with three-month improvement in behavioural responsiveness in patients with UWS. For the fMRI analyses (Fig. 1B), cortical and thalamic BOLD signals were transformed into edge time series and clustered into edge communities to characterize network organization using community similarity, normalized entropy and triadic motifs. Key thalamic nuclei identified by these analyses were then examined using NPI to estimate directed effective connectivity.

**Figure 1.**
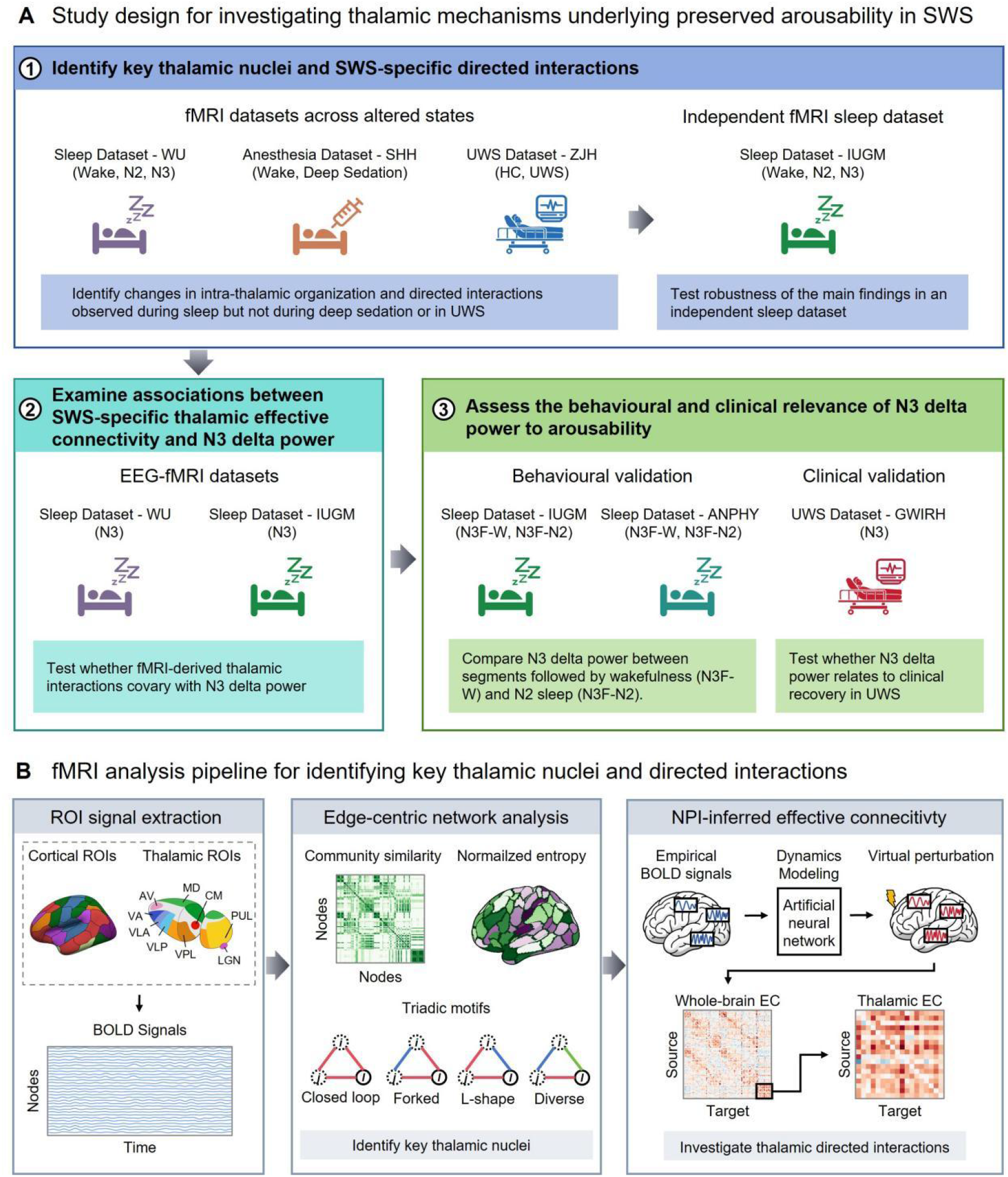
Study design and analytical workflow. (A) Study design for investigating thalamic mechanisms underlying preserved arousability in slow-wave sleep (SWS): (1) Sleep Dataset-WU, Anesthesia Dataset-SHH and UWS Dataset-ZJH served as the fMRI discovery datasets to identify changes in intra-thalamic organization and directed interactions observed during sleep but not during propofol-induced deep sedation or in unresponsive wakefulness syndrome (UWS). Sleep Dataset-IUGM was used as an independent fMRI replication dataset to assess the robustness of these findings. (2) Simultaneous EEG–fMRI data from Sleep Dataset-WU and Sleep Dataset-IUGM were used to examine whether the identified directed effective connectivity covaried with N3 delta power. (3) The behavioural relevance of N3 delta power was assessed by comparing N3 segments followed by wakefulness (N3F-W) with those followed by N2 sleep (N3F-N2) in Sleep Dataset-IUGM and Sleep Dataset-ANPHY. Its clinical relevance was evaluated by examining the association between N3 delta power and three-month improvement in behavioural responsiveness in UWS patients from UWS Dataset-GWIRH. (B) fMRI analysis pipeline for identifying key thalamic nuclei and directed interactions. BOLD signals were extracted from cortical and thalamic regions of interest (ROIs) and transformed into edge time series. Edge-centric analysis characterized overlapping network organization using community similarity, normalized entropy and triadic motifs to identify key thalamic nuclei. Neural perturbational inference (NPI) was then used to estimate directed effective connectivity (EC) among the identified thalamic nuclei through virtual perturbations of a neural-network surrogate trained on BOLD signals.

Here, we identified a CM/VPL-centred reconfiguration of intra-thalamic organization during N2 and N3 sleep, with selectively enhanced CM→VPL→PUL effective connectivity during N3. No analogous changes were observed during propofol-induced deep sedation or UWS. These directed interactions covaried with N3 delta power, which was associated with subsequent awakening and clinical recovery of behavioural responsiveness. Our findings highlight coordinated interactions among thalamic nuclei as a candidate neural mechanism supporting preserved arousability during slow-wave sleep.

## Results

We analysed six datasets spanning physiological sleep, propofol anesthesia and UWS (Fig. 1A; Table 1). Two simultaneous EEG-fMRI sleep datasets, Sleep Dataset-WU (Western University) and Sleep Dataset-IUGM (Institut Universitaire de Gériatrie de Montréal), included wakefulness, N2 and N3 sleep. Two additional fMRI datasets included Anesthesia Dataset-SHH (Shanghai Huashan Hospital), comprising wakefulness and propofol-induced deep sedation, and UWS Dataset-ZJH (Zhujiang Hospital), comprising healthy controls and patients with UWS. Two additional EEG datasets provided complementary validation: Sleep Dataset-ANPHY comprised overnight sleep recordings from healthy individuals, and UWS Dataset-GWIRH (Guangdong Work Injury Rehabilitation Hospital) comprised overnight EEG recordings from patients with UWS with three-month clinical follow-up. Participant demographic characteristics are summarized in Table 1.

**Table 1.** Demographic characteristics of participants across datasets. Sample sizes and demographic characteristics are reported separately for each analysis condition. Participants within the same dataset could contribute data to more than one condition. For the simultaneous EEG–fMRI datasets, EEG data temporally corresponding to the N3 fMRI segments were additionally used to assess associations between N3 delta power and fMRI-derived effective connectivity. Additional demographic and dataset information is provided in the Methods.

| Dataset | Analysis | Condition | Participants (n) | Sex (male) | Age (years) |
|---|---|---|---|---|---|
| **Simultaneous EEG-fMRI datasets** |  |  |  |  |  |
| Sleep Dataset-WU | fMRI analyses | W | 19 | 7 | 23.90 ± 3.76 |
|  |  | N2 | 25 | 10 | 24.36 ± 4.04 |
|  |  | N3 | 10 | 4 | 22.30 ± 4.06 |
| Sleep Dataset-IUGM | fMRI analyses | W | 11 | 5 | 27.09 ± 3.96 |
|  |  | N2 | 12 | 5 | 27.52 ± 3.91 |
|  |  | N3 | 8 | 4 | 26.25 ± 3.01 |
|  | N3 EEG awakening analysis | N3F-W | 5 | 3 | 27.00 ± 2.74 |
|  |  | N3F-N2 | 8 | 4 | 27.75 ± 3.99 |
| **fMRI datasets** |  |  |  |  |  |
| Anesthesia Dataset-SHH | fMRI analyses | W / DS | 24 | 12 | 46.33 ± 11.39 |
| UWS Dataset-ZJH | fMRI analyses | HC | 20 | 10 | 32.35 ± 10.69 |
|  |  | UWS | 20 | 11 | 51.60 ± 10.78 |
| **EEG datasets** |  |  |  |  |  |
| Sleep Dataset-ANPHY | N3 EEG awakening analysis | N3F-W | 19 | 8 | 31.74 ± 7.29 |
|  |  | N3F-N2 | 26 | 13 | 31.92 ± 6.03 |
| UWS Dataset-GWIRH | N3 EEG clinical recovery analysis | N3 | 10 | 7 | 53.10 ± 13.10 |

Sleep Dataset-WU, Anesthesia Dataset-SHH and UWS Dataset-ZJH first served as the fMRI discovery datasets to identify key thalamic nuclei showing changes in intra-thalamic organization and directed interactions during sleep, with no analogous changes observed during deep sedation or in UWS. Sleep Dataset-IUGM was then used to assess the reproducibility of these findings. Simultaneous EEG-fMRI data from Sleep Dataset-WU and Sleep Dataset-IUGM were used to examine whether the identified directed effective connectivity covaried with N3 delta power. Finally, Sleep Dataset-IUGM and Sleep Dataset-ANPHY were used to compare N3 delta power between N3F-W and N3F-N2, while UWS Dataset-GWIRH was used to test the association between N3 delta power and recovery of behavioural responsiveness in UWS patients. The overall study design and fMRI analysis pipeline are summarized in Fig. 1A and Fig. 1B, respectively.

### Edge-centric analyses reveal a distinctive intra-thalamic reconfiguration during N2 and N3 sleep

To characterize the overlapping organization of thalamic nuclei across altered states of consciousness, we applied edge-centric network analysis with cluster number = 10 to three datasets: Sleep Dataset-WU, Anesthesia Dataset-SHH and UWS Dataset-ZJH. We quantified three complementary properties of edge-community organization: community similarity, normalized entropy and triadic motif composition. These analyses revealed a CM/VPL-centered reconfiguration of intra-thalamic organization during N2 and N3 sleep, as detailed below.

### CM and VPL show enhanced convergence in edge-community profiles during N2 and N3 sleep

We first quantified community similarity, which measures the extent to which two nodes exhibit similar whole-brain edge-community profiles. For each thalamic nucleus, community similarity was averaged across its pairwise similarities with all other thalamic nuclei.

In Sleep Dataset-WU, linear mixed-effects model (LMM), with sleep stage as a fixed effect and subject as a random intercept, revealed a significant effect of sleep stage on community similarity in both CM and VPL (CM: F(2,51) = 6.791, p_Bonf_ = 0.022; VPL: F(2, 34.382) = 6.379, p_Bonf_ = 0.040) (Fig. 2, Table S1). Post hoc comparisons showed that community similarity was significantly higher in both N2 and N3 than in wakefulness for CM (W vs N2, estimate = −0.180, SE = 0.056, z = −3.199, p_Bonf_ = 0.004; W vs N3, estimate = −0.221, SE = 0.072, z = −3.057, p_Bonf_ = 0.007) and VPL (W vs N2, estimate = −0.155, SE = 0.050, z = −3.133, p_Bonf_ = 0.005; W vs N3, estimate = −0.194, SE = 0.066, z = −2.923, p_Bonf_ = 0.010). No significant difference was observed between N2 and N3 in either nucleus (CM: estimate = −0.041, SE = 0.069, z = −0.589, p_Bonf_ = 1.000; VPL: estimate = −0.038, SE = 0.063, z = −0.610, p_Bonf_ = 1.000; Fig. 2, Table S2).

**Figure 2.**
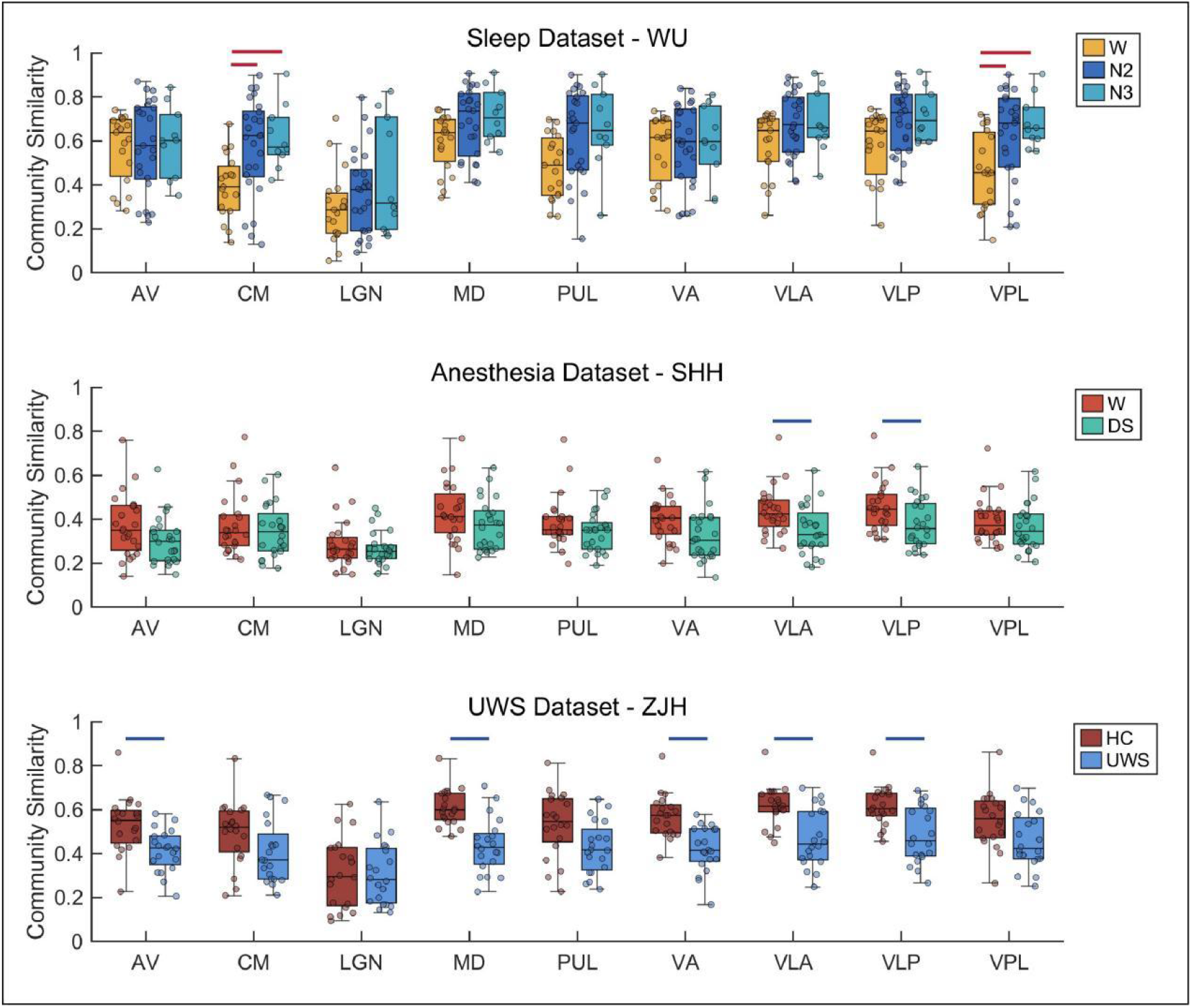
Comparison of thalamic community similarity across conditions in Sleep Dataset-WU, Anesthesia Dataset-SHH, and UWS Dataset-ZJH. Red line denotes significant increases and blue line denotes significant decreases at p < 0.05 (Bonferroni-corrected). Center line indicates median, box indicates interquartile range, and whisker denotes 1.5 × IQR. W = wakefulness; DS = deep sedation; HC = healthy controls; UWS = unresponsive wakefulness syndrome.

No corresponding increases in community similarity were observed in propofol-induced deep sedation or in UWS. Instead, community similarity was reduced in VLA and VLP during deep sedation relative to wakefulness and in AV, MD, VA, VLA and VLP in patients with UWS relative to healthy controls (all p_Bonf_ < 0.05; Fig. 2, Table S3-S4).

We next examined whether the increased similarity was specifically shown in CM and VPL rather than in cortical networks. Within-network community similarity was calculated by averaging all pairwise similarities among cortical regions belonging to each canonical network. No cortical network showed a corresponding increase during N2 or N3 sleep, deep sedation or UWS. Instead, community similarity was reduced in the dorsal attention network during sleep, with additional reductions observed across selected cortical networks during deep sedation and UWS (Fig. S1; Tables S5–S8).

Together, these findings identify a selective convergence of CM and VPL edge-community profiles during sleep, rather than a global increase in community similarity across cortical and thalamic systems.

### Thalamic edge-community diversity remains stable across states

We next examined normalized entropy, which quantifies how evenly the edges connected to each node are distributed across communities. No thalamic nucleus showed a significant change in normalized entropy across wakefulness, N2 and N3 sleep, between wakefulness and deep sedation, or between healthy controls and patients with UWS (Fig. S2; Tables S9 – S11). By contrast, several cortical networks exhibited state-dependent changes in normalized entropy. Entropy was reduced in the dorsal attention network during N2 and N3 sleep relative to wakefulness, increased in the frontoparietal and default-mode networks during deep sedation, and increased in the limbic, frontoparietal and default-mode networks in UWS (Fig. S3; Tables S12 – S15). Thus, the sleep-specific increases in CM and VPL community similarity reflect greater convergence of their edge-community profiles rather than changes in the diversity of their community participation.

### CM/VPL-centered triads shift towards closed-loop organization during N2 and N3 sleep

We next examined whether the sleep-specific convergence of CM and VPL was accompanied by changes in higher-order intra-thalamic organization. Triads were constructed using CM or VPL as the reference nucleus together with two other thalamic nuclei and were classified as closed-loop, forked, L-shape or diverse motifs according to the edge-community assignments of their constituent edges (Fig. 3A).

**Figure 3.**
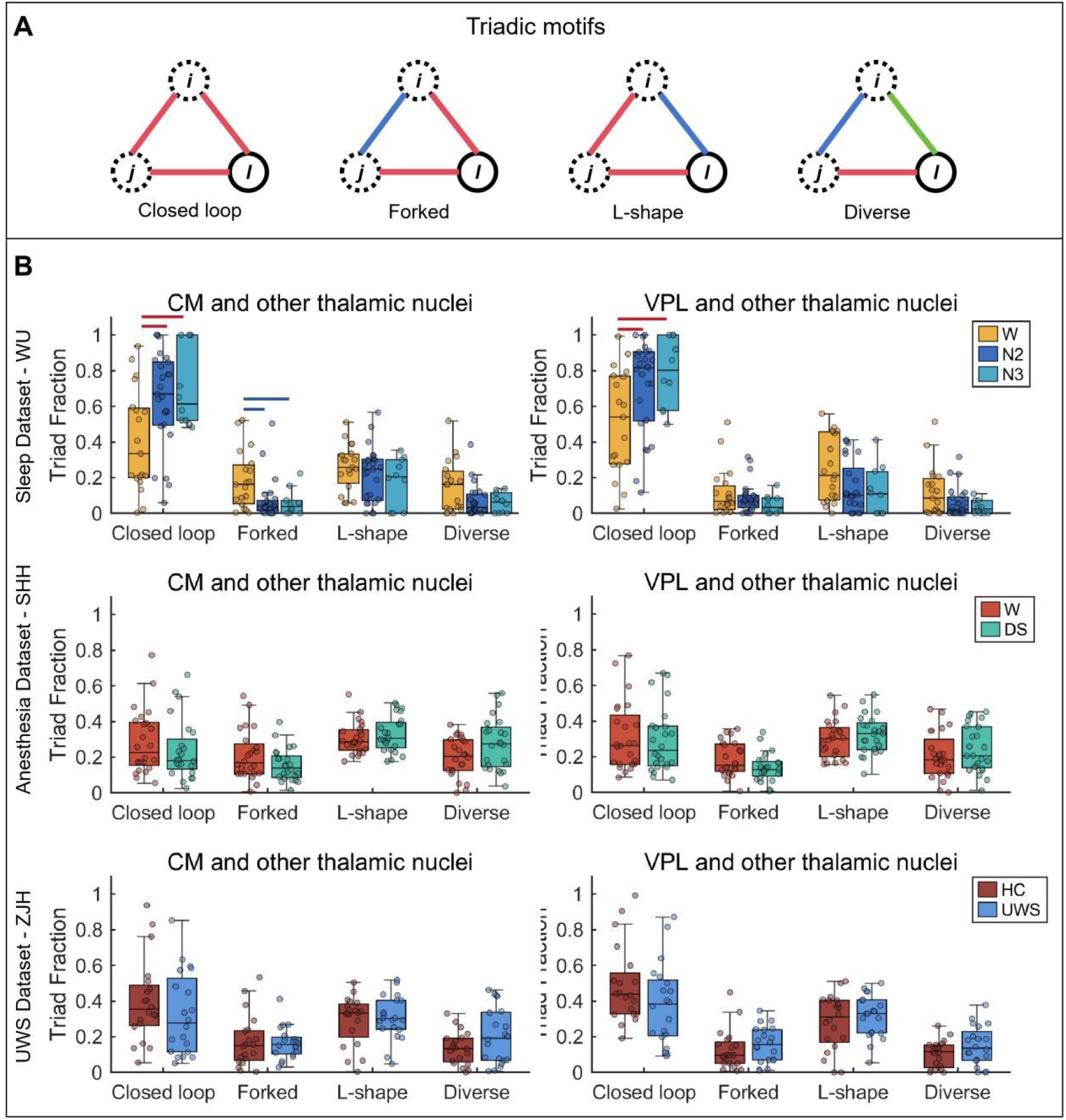
Thalamic-thalamic triadic motif organization centered on CM and VPL across Sleep Dataset-WU, Anesthesia Dataset-SHH, and UWS Dataset-ZJH. (A) Four types of triadic motifs defined for a reference node and the other two nodes: closed loop, forked, L-shape, and diverse. Edges shown in the same color indicate the same edge community assignment. (B) Comparison of the fraction of each motif type centered on CM and VPL across conditions in three datasets. Rows correspond to the datasets. Red line denotes significant increases and blue line denotes significant decreases at p < 0.05 (Bonferroni-corrected). Center line indicates median, box indicates interquartile range, and whisker denotes 1.5 × IQR. W = wakefulness; DS = deep sedation; HC = healthy controls; UWS = unresponsive wakefulness syndrome.

LMM showed significant effects of sleep stage on fraction of closed-loop motifs in both CM- and VPL-centered triads (CM: F(2,51) = 5.529, p_Bonf_ = 0.027; VPL: F(2, 36.168) = 5.173, p_Bonf_ = 0.042). Post hoc comparison showed that closed-loop motifs were more prevalent during both N2 and N3 than during wakefulness in CM-centered triads (W vs N2, estimate = −0.235, SE = 0.082, z = −2.876, p_Bonf_ = 0.012; W vs N3, estimate = −0.291, SE = 0.105, z = −2.770, p_Bonf_ = 0.017) and VPL-centered triads (W vs N2, estimate = −0.198, SE = 0.072, z = −2.774, p_Bonf_ = 0.017; W vs N3, estimate = −0.256, SE = 0.095, z = −2.687, p_Bonf_ = 0.022). No significant difference was observed between N2 and N3 in either nucleus (Fig. 3B, Table S16-S17).

Sleep stage also significantly affected the fraction of forked motifs in CM-centered triads (F(2,51) = 5.129, p_Bonf_ = 0.037). Forked motifs were less prevalent during both N2 and N3 than during wakefulness (W vs N2: estimate = 0.113, SE = 0.039, z = 2.895, p_Bonf_ = 0.011; W vs N3: estimate = 0.126, SE = 0.050, z = 2.511, p_Bonf_ = 0.036), with no significant difference between N2 and N3 (Fig. 3B, Table S16-S17). In addition, no corresponding changes in thalamic-thalamic motif composition were observed during deep sedation or in UWS (Fig. 3B; Tables S18 – S19). These findings indicate that N2 and N3 sleep are characterized by a shift towards more coherent CM/VPL-centered organization within the thalamus.

As complementary analyses, we examined thalamo-cortical triads by taking CM or VPL as the reference node and pairing it with two cortical regions. Sleep stage also significantly affected the fraction of closed-loop motifs in both CM- and VPL-centered thalamo-cortical triads (CM: F(2,36.387) = 7.871, p_Bonf_ = 0.006; VPL: F(2, 32.546) = 5.488, p_Bonf_ = 0.035). However, the direction of change was opposite to that observed in intra-thalamic triads: closed-loop motifs were less prevalent during both N2 and N3 than during wakefulness in CM-centered triads (W vs N2, estimate = 0.029, SE = 0.009, z = 3.093, p_Bonf_ = 0.006; W vs N3, estimate = 0.045, SE = 0.012, z = 3.596, p_Bonf_ = 0.001) and VPL-centered triads (W vs N2, estimate = 0.020, SE = 0.008, z = 2.432, p_Bonf_ = 0.045; W vs N3, estimate = 0.035, SE = 0.011, z = 3.093, p_Bonf_ = 0.006). No significant difference was observed between N2 and N3 in either nucleus (Fig. S4, Table S20-S21). Moreover, no thalamo-cortical motif type differed significantly between wakefulness and deep sedation or between healthy controls and patients with UWS, except for a reduced fraction of L-shape motifs in VPL-centered triads during deep sedation (t(23) = 2.86, p_Bonf_ = 0.035; Fig. S4 and Table S22-S23).

Together, these contrasting patterns show that sleep increases coherent CM/VPL-centered triadic organization within the thalamus while reducing coherent CM/VPL-centered organization across thalamo-cortical configurations.

### Edge-centric findings are robust to the number of edge communities

Because edge-centric analysis depends on the choice of the number of edge communities, the primary analyses were conducted using = 10, consistent with previous edge-centric studies ^56,67,68^. To assess robustness, we repeated all edge-centric analyses across cluster number of from 3 to 20 (excluding = 10). The principal findings for community similarity, normalized entropy and triadic motif composition were reproduced across multiple cluster numbers, particularly = 11-13 and 16-20 (Fig. S5–S10; Data S1–S3), indicating that the results were robust across clustering number.

### Directed CM→VPL→PUL interactions are selectively enhanced during N3 sleep

Building on the edge-centric findings that identified CM and VPL as key nuclei in the sleep-specific reconfiguration of intra-thalamic organization, and on prior evidence that CM activation precedes VPL and is followed by PUL during the sleep-to-wake transition^21^, we next asked whether this candidate CM→VPL→PUL sequence showed enhanced directed interactions during sleep (Fig. 4A-B). Neural perturbational inference was used to estimate effective connectivity from CM to VPL and from VPL to PUL. Because NPI depends on virtual perturbations of a trained surrogate model, we computed effective connectivity across a range of perturbing strength (1–10, step = 1) and averaged the resulting connectivity to reduce sensitivity to a specific strength choice.

**Figure 4.**
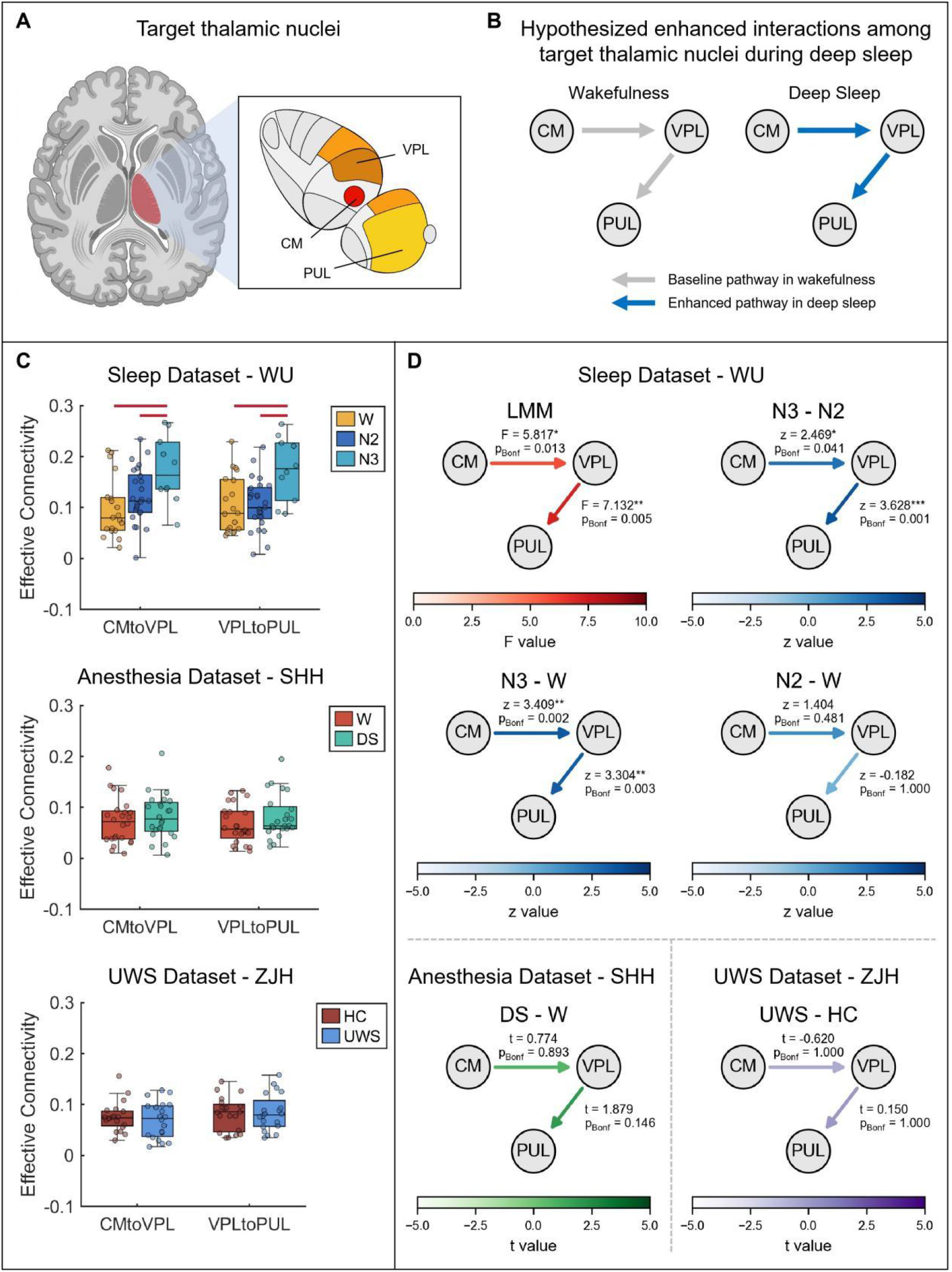
Inferred effective connectivity among target thalamic nuclei. (A) Schematic illustration of target thalamic nuclei identified in edge-centric analysis, including CM, VPL and PUL. (B) Hypothesized enhanced interactions among target thalamic nuclei during deep sleep. (C) Comparison of effective connectivity for CM→VPL and VPL→PUL across conditions in Sleep Dataset-WU, Anesthesia Dataset-SHH, and UWS Dataset-ZJH. Red line denotes significant increases at p < 0.05 (Bonferroni-corrected). Center line indicates median, box indicates interquartile range, and whisker denotes 1.5 × IQR. (D) Statistical summary of condition effects for these directed connections, including overall linear mixed-effects model results (F values) and post hoc contrasts (z values) in the Sleep Dataset-WU, and between-condition comparisons (t values) in the Anesthesia Dataset-SHH and UWS Dataset-ZJH. W = wakefulness; DS = deep sedation; HC = healthy controls; UWS = unresponsive wakefulness syndrome.

LMM revealed significant effects of sleep stage on both CM→VPL and VPL→PUL effective connectivity (CM→VPL: F(2, 34.715) = 5.817, p_Bonf_ = 0.013; VPL→PUL: F(2, 34.735) = 7.132, p_Bonf_ = 0.005; Fig. 4C, D, Table S24). Post hoc comparisons showed that both connections were stronger during N3 than during wakefulness and N2 sleep. CM→VPL effective connectivity was higher in N3 than in wakefulness (estimate = −0.074, SE = 0.022, z = −3.409, p_Bonf_ = 0.002) and N2 (estimate = −0.051, SE = 0.021, z = −2.469, p_Bonf_ = 0.041). Similarly, VPL → PUL effective connectivity was higher in N3 than in wakefulness (estimate = −0.066, SE = 0.020, z = −3.304, p_Bonf_ = 0.003) and N2 (estimate = −0.068, SE = 0.019, z = −3.628, p_Bonf_ = 0.001). Neither connection differed significantly between wakefulness and N2 (Fig. 4C, D and Table S25). Moreover, neither CM→VPL nor VPL→PUL effective connectivity differed significantly between wakefulness and deep sedation or between healthy controls and patients with UWS (Fig. 4C,D and Table S26–S27). Together, these findings indicate that directed interactions along the CM→VPL→PUL sequence were selectively strengthened during N3, rather than being a general feature of NREM sleep, with no analogous enhancement observed in deep sedation or UWS.

To assess how well the surrogate model reproduced the empirical BOLD signals, we compared state-averaged functional connectivity matrices derived from model-generated and empirical BOLD signals. Across perturbation strengths from 1 to 10, model-derived functional connectivity was strongly correlated with empirical functional connectivity in every state (r = 0.94 – 0.98, all p < 0.001; Fig. S11 – S16), indicating that the surrogate models reproduced the large-scale inter-regional correlation structure of the empirical data.

### Independent replication of sleep-specific thalamic coordination

We next assessed the reproducibility of the sleep-related thalamic coordination pattern in an independent sleep dataset (Sleep Dataset-IUGM) using the same edge-centric (k = 10) and NPI pipelines. Replication analyses focused on the a priori features identified in Sleep Dataset-WU: CM and VPL community similarity, the fraction of closed-loop motifs in CM- and VPL-centered intra-thalamic triads, and effective connectivity from CM to VPL and from VPL to PUL.

Community similarity findings were robustly replicated. Linear mixed-effects models revealed significant effects of sleep stage in both CM and VPL (CM: F(2,17.923) = 10.060, p_Bonf_ = 0.002; VPL: F(2, 17.416) = 7.666, p_Bonf_ = 0.008; Fig. S17A, Table S28). Post hoc comparison showed that community similarity was higher during both N2 and N3 than during wakefulness in CM (W vs N2, estimate = −0.273, SE = 0.071, z = −3.845, p_Bonf_ < 0.001; W vs N3, estimate = −0.298, SE = 0.079, z = −3.782, p_Bonf_ < 0.001) and VPL (W vs N2, estimate = −0.176, SE = 0.066, z = −2.658, p_Bonf_ = 0.024; W vs N3, estimate = −0.277, SE = 0.074, z = −3.763, p_Bonf_ = 0.001). Neither nucleus differed significantly between N2 and N3 (Fig. S17A and Table S29).

Evidence for replication of the closed-loop motif findings was limited. The overall effect of sleep stage did not remain significant after correction in either CM- or VPL-centered triads (CM: F(2,17.019) = 3.767, p_uncorrect_ = 0.044, p_Bonf_ = 0.177; VPL: F(2, 18.071) = 2.398, p_uncorrect_ = 0.119, p_Bonf_ = 0.477; Fig. S17B, Table S30). However, pairwise comparison showed that the fraction of closed-loop motifs was significantly higher in N3 than in wakefulness for CM-centered triads after correction (estimate = −0.339, SE = 0.126, z = −2.698, p_uncorrect_ = 0.007, p_Bonf_ = 0.021). The corresponding difference in VPL-centered triads was in the same direction but did not remain significant after correction (estimate = −0.254, SE = 0.124, z = −2.041, p_uncorrect_ = 0.041, p_Bonf_ = 0.124; Fig. S17B, Table S31). In the NPI analysis, the overall effects of sleep stage were not significant for either CM→ VPL or VPL→PUL effective connectivity (CM→VPL: F(2,21.037) = 1.474, p_uncorrect_ = 0.252, p_Bonf_ = 0.504; VPL→PUL: F(2, 20.464) = 2.265, p_uncorrect_ = 0.129, p_Bonf_ = 0.259; Fig. S17C, Table S32).

Nevertheless, the estimated effects were consistently aligned with those observed in Sleep Dataset-WU. CM→VPL effective connectivity was higher in N3 than in wakefulness (estimate = −0.023, SE = 0.032, z = −0.727, p_uncorrect_ = 0.467) and N2 (estimate = −0.053, SE = 0.031, z = −1.684, p_uncorrect_ = 0.092). Similarly, VPL → PUL effective connectivity was higher in N3 than in wakefulness (estimate = −0.049, SE = 0.028, z = −1.777, p_uncorrect_ = 0.075), and N2 (estimate = −0.054, SE = 0.027, z = −1.997, p_uncorrect_ = 0.046). Notably, the N3 vs. N2 difference in VPL→PUL effective connectivity reached significance at the uncorrected level, although it did not survive Bonferroni correction (p_Bonf_ = 0.138); none of the remaining pairwise comparisons was significant after correction (Fig. S17C and Table S33). Model-derived functional connectivity was also strongly correlated with empirical functional connectivity across all states and perturbation strengths (r = 0.95 − 0.97, all p < 0.001; Fig. S18–S19).

Overall, the independent dataset robustly replicated the increase in CM and VPL community similarity and provided limited support for increased closed-loop organization. The NPI-derived effects did not survive correction, possibly owing to the smaller N3 sample, but their directions were consistent with the discovery findings, providing supportive rather than confirmatory evidence for N3-specific enhancement of the CM→VPL→PUL effective connectivity.

### N3 delta power links directed intra-thalamic interactions to awakening propensity and clinical recovery

The selective enhancement of CM → VPL → PUL effective connectivity during N3 sleep identified a candidate thalamic pathway for preserved arousability. However, its N3 specificity alone was insufficient to establish its relevance to arousability. We therefore hypothesized that, if this pathway contributes to arousability during N3 sleep, its strength should be associated with an electrophysiological feature of N3 that is also related to subsequent awakening. Because delta activity is a defining feature of N3 sleep and is closely linked to thalamic activity^44,63–65^, we first tested whether relative delta power covaried with CM → VPL and VPL → PUL effective connectivity. We then examined whether N3 delta power was associated with subsequent awakening in healthy individuals and recovery of behavioural responsiveness in patients with UWS.

To test the first hypothesis, we pooled data from Sleep Dataset-WU and Sleep Dataset-IUGM and examined the associations between relative delta power and fMRI-derived effective connectivity. Partial Pearson correlations were performed while controlling for dataset identity. Relative delta power (1-4 Hz) was strongly and positively associated with both CM → VPL effective connectivity (r = 0.611, p_Bonf_ = 0.018) and VPL→PUL effective connectivity (r = 0.716, p_Bonf_ = 0.002) during N3 sleep (Fig. 5A). These findings identify cortical delta power as an electrophysiological correlate of enhanced directed intra-thalamic interactions during N3 sleep.

**Figure 5.**
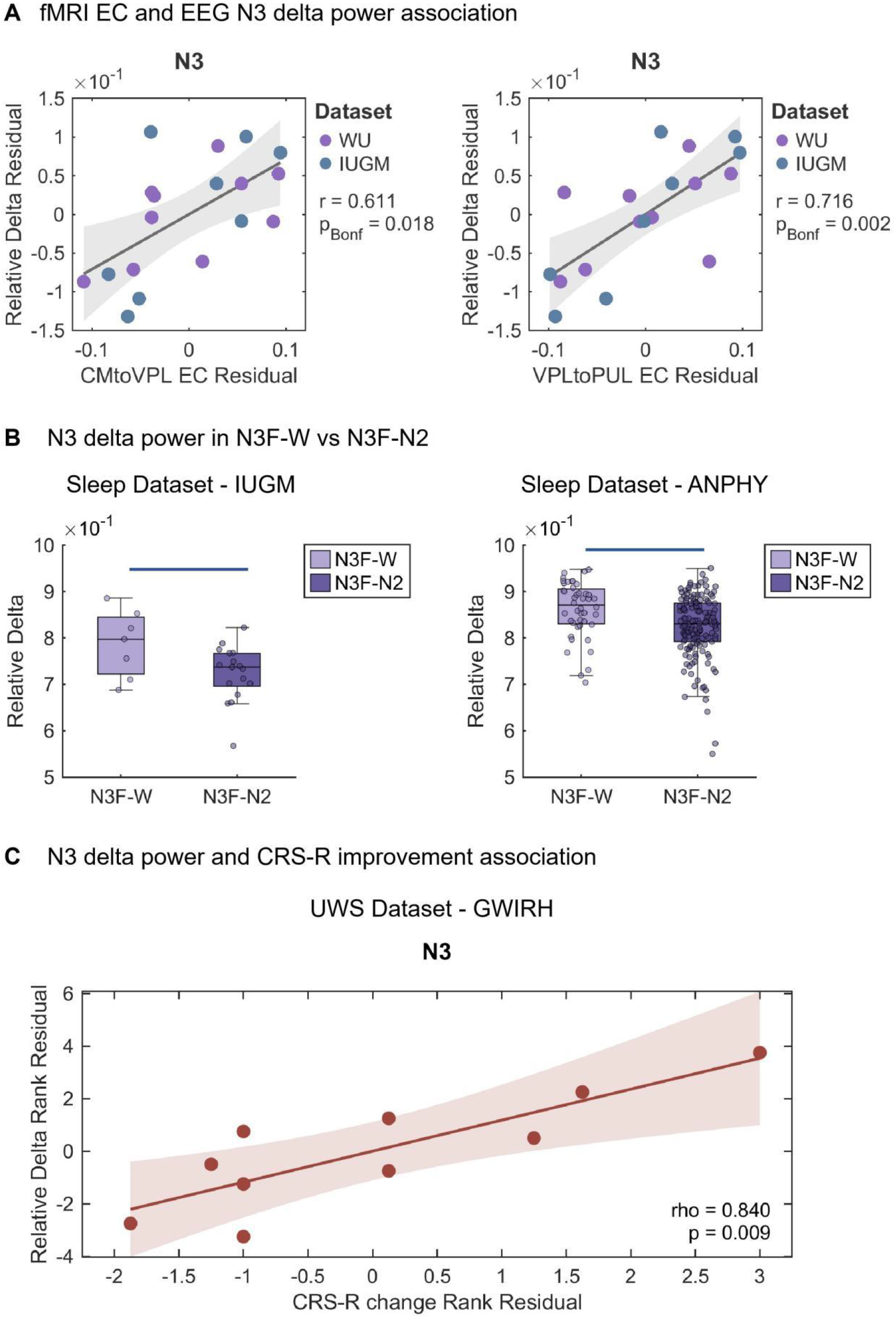
N3 delta power links directed intra-thalamic interactions to awakening propensity and clinical recovery. (A) Associations between N3 relative delta power and fMRI-derived effective connectivity (EC) along the CM→VPL→PUL pathway. Data from Sleep Dataset-WU and Sleep Dataset-IUGM were pooled, and partial Pearson correlations were performed while controlling for dataset identity. Relative delta power was positively associated with both CM→VPL EC (r = 0.611, p_Bonf_ = 0.018) and VPL→PUL EC (r = 0.716, p_Bonf_ = 0.002). (B) N3 relative delta power in N3 segments followed by wakefulness (N3F-W) or N2 sleep (N3F-N2) in Sleep Dataset-IUGM (N3F-W, n = 7 segments; N3F-N2, n = 17 segments) and Sleep Dataset-ANPHY (N3F-W, n = 44 segments; N3F-N2, n = 148 segments). Relative delta power was quantified over the final 60 s of each N3 segment and compared at the segment level using linear mixed-effects models, with subsequent state as a fixed effect and participant as a random intercept. Relative delta power was significantly higher in N3F-W than in N3F-N2 in both datasets (Sleep Dataset-IUGM: estimate = 0.063, SE = 0.024, t = 2.632, p = 0.008; Sleep Dataset-ANPHY: estimate = 0.040, SE = 0.011, t = 3.728, p < 0.001; one-tailed tests). (C) Association between baseline N3 relative delta power and subsequent improvement in behavioural responsiveness in patients with UWS from UWS Dataset-GWIRH. Clinical improvement was quantified as the change in CRS-R score from baseline to the three-month follow-up. Partial Spearman correlation, controlling for injury etiology, showed a positive association between baseline N3 relative delta power and CRS-R improvement (rho = 0.840, p = 0.009; two-tailed).

We next asked whether N3 delta power was related to the propensity for imminent awakening. We therefore compared relative delta power between N3F-W and N3F-N2 in Sleep Dataset-IUGM and Sleep Dataset-ANPHY. Sleep Dataset-WU was excluded because it contained only one N3F-W segment. Relative delta power was quantified over the final 60 s of each N3 segment and analysed at the segment level using linear mixed-effects models, with subsequent state as a fixed effect and participant as a random intercept. Based on the hypothesis that greater N3 delta power would be associated with a higher propensity to awaken, one-tailed tests were used. In both datasets, relative delta power was significantly higher in N3F-W than in N3F-N2 segments (Sleep Dataset-IUGM: estimate = 0.063, SE = 0.024, t = 2.632, p = 0.008; Sleep Dataset-ANPHY: estimate = 0.040, SE = 0.011, t = 3.728, p < 0.001; one-tailed tests; Fig. 5B). Thus, greater relative delta power during N3 was associated with subsequent transitions to wakefulness rather than N2 sleep.

Finally, we examined whether N3 delta power was associated with recovery of behavioural responsiveness in patients with UWS. In UWS Dataset-GWIRH, clinical improvement was quantified as the change in CRS-R score from baseline to the three-month follow-up. After controlling for injury aetiology, partial Spearman correlation revealed a strong positive association between N3 relative delta power and CRS-R improvement in UWS patients (rho = 0.84, p = 0.009, two-tailed; Fig. 5C). This result extends the relevance of N3 delta power from physiological sleep to a clinical population, linking greater delta activity during N3 sleep to subsequent recovery of behavioural responsiveness in UWS patients.

Together, these cross-modal findings provide empirical support for the proposed involvement of the CM→VPL→PUL effective connectivity in sleep-specific arousability. The strength of both directed connections covaried with relative delta power during N3 sleep, while N3 delta power was associated with a greater propensity to transition from N3 sleep to wakefulness in healthy individuals and with greater recovery of behavioural responsiveness in patients with UWS. Thus, convergent physiological, behavioural and clinical evidence supports enhanced CM→VPL→PUL effective connectivity as a candidate thalamic mechanism for preserved arousability during N3 sleep.

## Discussion

Our study identifies that a CM–VPL–PUL-centred pattern of intra-thalamic coordination is associated with preserved arousability during sleep. Edge-centric analyses revealed increased CM/VPL community similarity and a shift towards closed-loop intra-thalamic organization during both N2 and N3 sleep, whereas NPI showed selectively enhanced directed effective connectivity along the CM→VPL→PUL pathway during N3. These sleep-related changes were not observed during propofol-induced deep sedation or in UWS. The increase in CM/VPL community similarity was robustly replicated in an independent sleep dataset, with limited support for the motif and effective-connectivity findings. Moreover, CM → VPL and VPL → PUL effective connectivity covaried with N3 delta power, which was associated with a greater propensity for subsequent awakening in healthy individuals and greater recovery of behavioural responsiveness in patients with UWS. Together, these findings jointly demonstrate that the CM-VPL-PUL-centred thalamic coordination functions as a candidate mechanism underlying preserved arousability during sleep (Fig. 6).

**Figure 6.**
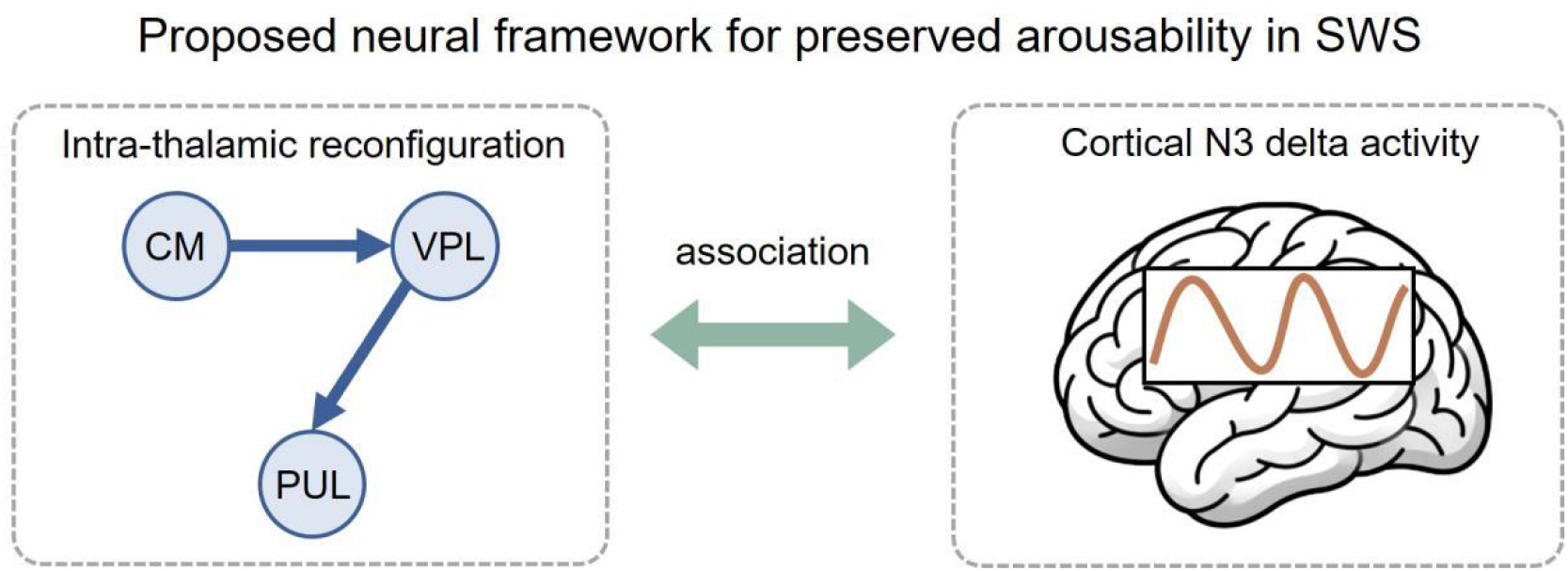
Proposed neural framework for preserved arousability in sleep. The schematic highlights selectively enhanced effective connectivity along the CM→VPL→PUL pathway during N3 sleep as a candidate intra-thalamic mechanism supporting preserved arousability. The strength of CM→VPL and VPL→PUL effective connectivity was positively associated with cortical delta activity during N3, while N3 delta power was further associated with subsequent awakening in healthy individuals and recovery of behavioural responsiveness in patients with UWS. These findings suggest that coordinated interactions along the CM→VPL→PUL pathway may contribute to preserved arousability during slow-wave sleep.

During N2 and N3 sleep, thalamic functional architecture was reorganized towards a CM/VPL-centred coordination pattern. In the edge-centric framework, increased community similarity indicates that CM and VPL share more similar edge-community profiles across their connections with other thalamic nuclei, reflecting greater convergence within overlapping intra-thalamic subnetworks. This convergence was accompanied by an increased prevalence of CM/VPL-centred closed-loop motifs, indicating that CM/VPL and other thalamic nuclei more frequently coordinated through the same edge community. Furthermore, these changes occurred without alterations in thalamic normalized entropy, suggesting that the reconfiguration reflected convergence of community profiles rather than a general reduction in the diversity of community participation. Additionally, no cortical network showed a corresponding increase in community similarity, while CM/VPL-centred closed-loop motifs decreased in thalamo-cortical configurations during N2 and N3. Together, these complementary findings suggest a selective shift towards more coherent intra-thalamic organization during N2 and N3 sleep rather than a global increase in network coherence. This interpretation is consistent with previous evidence that thalamic activity is recruited during arousal events in sleep^69^ and that intra-thalamic functional coupling increases during N2 sleep^70^. Moreover, CM receives input from the ascending reticular activating system^71–74^ and has been directly linked to behavioural arousal through stimulation studies,^44,45,49–51,75^, whereas VPL co-activation with global brain activity has also been associated with arousal^5^. Thus, the CM/VPL-centred reconfiguration identified here may provide an intra-thalamic network architecture that helps preserve the capacity for awakening during sleep.

Our findings suggest two levels of intra-thalamic reconfiguration across N2 and N3 sleep. A CM/VPL-centred architectural reconfiguration was shared by N2 and N3, whereas directed effective connectivity along the CM→VPL→PUL pathway was selectively strengthened during N3, with both connections stronger than in wakefulness and N2 and no corresponding increase in N2 relative to wakefulness. Thus, N2 may provide an intermediate state, indicating that enhanced CM→VPL→PUL interaction is not uniformly observed across N2 and N3 sleep, but is selectively associated with N3. This distinction suggests that N2 and N3 share a broader intra-thalamic architectural shift, whereas N3 additionally engages stronger directed interactions along the CM→ VPL → PUL pathway. This N3-specific pattern is supported by the previous evidence of a temporally ordered CM → VPL → PUL activation sequence preceding awakening from NREM sleep^21^. CM, VPL and PUL have also each been linked to arousal-related processes, with evidence for CM and VPL discussed above and PUL implicated in arousal-related global co-activation^5^ and spontaneous arousal fluctuations^76^. Mechanistically, CM may relay arousal-related signals from the ascending reticular activating system^71–74^ to VPL, a major somatosensory relay^77,78^, and subsequently to PUL, a higher-order nucleus involved in sensory integration and visual attention^79–82^. Enhanced directed coordination along this pathway may therefore represent an additional thalamic mechanism for preserving awakening capacity during N3 sleep.

The association between N3 delta activity and CM→VPL→PUL effective connectivity provides an electrophysiological link between this directed thalamic pathway and arousability. Greater relative delta power during N3 covaried with stronger CM→VPL and VPL→PUL effective connectivity, suggesting that delta activity may reflect neural processes accompanying enhanced intra-thalamic coordination. Although delta activity is traditionally associated with sleep depth and diminished consciousness, prominent delta oscillations can also occur in conscious states, indicating that elevated delta activity does not necessarily signify reduced consciousness^23,66^. Consistent with this view, our findings showed that delta power was higher in N3 periods followed by wakefulness than in those followed by N2 sleep. A recent high-density EEG study similarly showed transient increases in delta activity in the seconds preceding arousal or awakening from NREM sleep, followed by high-frequency activation several seconds later^18^. Such arousal-related slow activity has also been proposed to reflect transient activation of subcortical arousal systems, including intralaminar thalamic nuclei, interacting with the bistable cortical dynamics of NREM sleep^18^. Thus, our findings suggest that, within N3, delta activity may also carry information about the preserved propensity for awakening. The clinical relevance of this interpretation was further supported by the positive association between baseline N3 delta power and subsequent improvement in behavioural responsiveness in UWS. Consistent with this finding, previous studies reported restoration of slow-wave regulation alongside clinical recovery in DOC^83^ and higher resting-state delta power in patients transitioning from UWS to MCS^84^. Together, these convergent physiological, behavioural and clinical findings broaden the functional significance of N3 delta activity beyond a marker of sleep depth, suggesting that it may index preserved arousability and provide indirect support for a link between intra-thalamic coordination and this capacity. We further hypothesize that, as sleep deepens and delta activity increases, a compensatory arousal-promoting mechanism may be recruited beyond a certain level of delta activity, potentially involving enhanced CM→VPL→PUL effective connectivity to facilitate awakening. Such a mechanism may help preserve the reversibility of deep sleep, although its threshold dependence and causal role remain to be tested.

The absence of analogous CM/VPL-centred reconfiguration and CM → VPL → PUL enhancement during propofol-induced deep sedation and UWS suggests that these changes are not a general consequence of reduced consciousness. Although propofol anesthesia and UWS arise from fundamentally different mechanisms, both are associated with disruption of thalamic function. Propofol potentiates GABA_A_ receptor-mediated inhibition, suppressing activity in thalamic nuclei including CM^42,43,85^ and VPL^42,86^ and disrupting thalamocortical interactions^30,87^. In UWS, the mesocircuit model proposes that reduced striatal output increases inhibitory pallidal influence on CM^52^. Neuroimaging studies further demonstrated thalamic metabolic dysfunction, reduced thalamic BOLD activity and impaired intrathalamic structural integrity^88–94^. These distinct forms of thalamic dysfunction may constrain the coordinated intra-thalamic dynamics observed during physiological sleep. By contrast, physiological sleep is an endogenous and readily reversible state in which the capacity for spontaneous or stimulus-evoked awakening is preserved. The sleep-specific intra-thalamic reconfiguration identified here may therefore reflect a thalamic organization compatible with this reversibility and preserved arousability, rather than reduced consciousness itself. Thalamic reticular regulation and reciprocal corticothalamic interactions may contribute to this sleep-specific thalamic coordination^65,95,96^ and warrant electrophysiological investigation.

Several limitations should be considered. First, although NPI has been validated by its ability to reproduce stimulation-evoked propagation patterns^59^, it remains a model-based approach for estimating directed effective connectivity from fMRI data. Further validation using empirical perturbation or electrophysiological recordings will help clarify the neural interactions among CM, VPL and PUL during sleep. Second, although the increase in CM/VPL community similarity was robustly replicated in the independent sleep dataset, support for the closed-loop motif and NPI findings was more limited, possibly owing in part to the smaller N3 sample. Larger sleep EEG– fMRI cohorts are therefore needed to confirm these higher-order and directed interaction patterns. Third, the link between CM→VPL→PUL effective connectivity and arousability is supported by its covariation with N3 delta power, which was also related to subsequent awakening and clinical recovery. Whether CM→VPL→PUL effective connectivity itself predicts awakening or recovery remains to be tested directly. Finally, the clinical association of N3 delta activity was derived from a relatively small UWS cohort and requires independent longitudinal replication in larger samples to establish its generalizability.

## Conclusion

In conclusion, our study identifies a sleep-specific intra-thalamic reconfiguration associated with preserved arousability. Edge-centric analyses revealed a CM/VPL-centred architectural reconfiguration shared by N2 and N3 sleep, and NPI further identified selectively enhanced directed effective connectivity along the CM→VPL →PUL pathway during N3. These changes were not observed during propofol-induced deep sedation or UWS, suggesting that they are not a general consequence of reduced consciousness. Moreover, CM → VPL → PUL effective connectivity covaried with N3 delta power, with the N3 delta power being related to subsequent awakening in healthy individuals and recovery of behavioural responsiveness in UWS. Together, these findings highlight coordinated interactions among thalamic nuclei as a candidate mechanism for preserving the capacity for awakening during slow-wave sleep and provide a complementary framework for understanding arousability beyond isolated thalamic nuclei or thalamocortical interactions.

## Reporting summary

Further information on research design is available in the Nature Portfolio Reporting Summary linked to this article.

## Data availability

The Sleep Dataset - ANPHY can be accessed at https://osf.io/r26fh/overview. Other fMRI and EEG datasets are available from the corresponding author upon reasonable request and subject to institutional approval.

## Code availability

The code for edge-centric network analysis is available at https://github.com/brain-networks/edge-centric_demo. The code for NPI are availabel at https://github.com/ncclab-sustech/NPI/.

## Acknowledgements

We gratefully acknowledge Stuart Fogel for sharing the sleep EEG– fMRI datasets used in this study. This work was supported by grants from the National Key Research and Development Program of China (No. 2024YFC3606900, No. 2025YFE0213500), National Natural Science Foundation of China (No. 32400861, No. 32371098), Natural Science Foundation of Guangdong Province (No. 2024A1515010527, No. 2024A1515011429).

## Author contributions

J.H. and P.Q. conceived and designed the study. J.H. performed the analyses, prepared the figures, and drafted and revised the manuscript. Q.X. designed the experiments and collected and curated the fMRI data from patients with UWS. J.Z. designed the experiments and collected and curated the anesthesia fMRI data. M.L., Y.L. designed the experiments and collected and curated the sleep EEG data from patients with UWS. Y.H., X.H., G.N., S.H. and P.Q. contributed to the interpretation of the results and reviewed the manuscript. P.Q. supervised the study. All authors reviewed and approved the final manuscript.

## Methods

### Participants and data acquisition

#### Simultaneous EEG-fMRI datasets: Sleep Dataset-WU

This dataset served as the basis for published studies ^5,97^. Thirty-five healthy right-handed adults (14 male, age 23.69 ± 3.62 years) passed the inclusion/exclusion criteria and slept in the scanner with a simultaneous EEG-fMRI recording. For each participant, up to 2 h of sleep EEG–fMRI data was acquired. EEG recordings obtained during simultaneous EEG–fMRI were scored by an expert registered polysomnographic technologist according to standard American Academy of Sleep Medicine (AASM) criteria.

For the fMRI analyses, the longest continuous segments of each sleep stage were selected to minimize discontinuities in data analysis. Participants with insufficient data for a given stage (< 90 fMRI volumes) were excluded from that stage-specific analysis, resulting in 19 participants for wakefulness, 25 for N2 sleep, and 10 for N3 sleep (Table 1). EEG data temporally corresponding to the retained N3 fMRI segments were additionally used for the EEG-fMRI association analysis described below. All study procedures and methods adhered to the Declaration of Helsinki and were approved by the Western University Health Science research ethics board. Additional details regarding participant eligibility and data acquisition are provided in the previous studies ^97^.

Functional magnetic resonance imaging was performed at a 3.0T Magnetom Prisma MR imaging system (Siemens, Erlangen, Germany) using a 64-channel head coil. High-resolution anatomic images were acquired using a standard 3D Multislice MPRAGE sequence (TR = 2300 ms, TE = 2.98 ms, TI = 900 ms, FA = 9^◦^, 176 slices, FoV = 256 × 256 mm^2^, matrix size = 256 × 256 × 176, voxel size = 1 × 1 × 1 mm^3^). During the sleep session, T2^∗^-weighted fMRI images were acquired with a gradient echo-planar imaging (EPI) sequence using axial slice orientation (TR = 2160 ms, TE = 30 ms, FA = 90^◦^, 40 transverse slices, 3 mm slice thickness, 10% inter-slice gap, FoV = 220 × 220 mm^2^, matrix size = 64 × 64 × 40, voxel size = 3.44 × 3.44 × 3 mm^3^).

Simultaneous EEG with MRI was recorded using a 64-channel MR-compatible cap with one ECG lead (BrainCap MR; Easycap, Herrsching, Germany) and two 32-channel MR-compatible amplifiers (BrainAmp MR plus; Brain Products GmbH, Gilching, Germany). EEG signals were referenced to FCz, and electrode impedances were maintained below 5 kΩ. To improve R-peak detection for ballistocardiographic (BCG) artifact correction, three additional bipolar ECG derivations were recorded using a 16-channel MR-compatible amplifier (BrainAmp ExG MR; Brain Products). Data were sampled at 5 kHz with a resolution of 500 nV/bit and analog-filtered using a 500-Hz low-pass filter and a 0.0159-Hz high-pass filter. Further details of simultaneous EEG–fMRI acquisition have been reported previously ^97^.

#### Simultaneous EEG-fMRI datasets: Sleep Dataset-IUGM

This dataset served as the basis for a published study ^98^. A total of 13 healthy right-handed adults (6 male, age 27.38 ± 3.78 years) met the inclusion/exclusion criteria and underwent simultaneous EEG–fMRI recording while sleeping in the scanner. The sleep session was terminated when the maximum possible number of volumes for a single fMRI session (4000 volumes, lasting a maximum of 2.5 hr) was reached, or if subjects voluntarily terminated the session. Wakefulness and NREM sleep stages were scored from EEG recordings according to standard AASM criteria.

For the fMRI analyses, the longest continuous segments of sleep stages were selected to minimize discontinuities in our data analysis. Participants with insufficient data for a given stage (< 90 fMRI volumes) were excluded from that stage-specific analysis, resulting in 11 participants for wakefulness, 12 for N2 sleep, and 8 for N3 sleep (Table 1). EEG data temporally corresponding to the retained N3 fMRI segments were additionally used for the EEG-fMRI association analysis described below. For the N3 EEG awakening analysis, 60-s N3 segments were further categorized according to the subsequent sleep state as N3F-W or N3F-N2. Five participants contributed N3F-W segments and eight contributed N3F-N2 segments (Table 1). Details of the EEG segment selection and spectral analyses are described below. Ethical and scientific approval was obtained from the Research Ethics Board at the IUGM, Montreal, Quebec, Canada, and written informed consent was obtained from all participants prior to the study. Additional details are provided in the previous study ^98^.

Images were collected using a 3T TIM TRIO Siemens scanner with a 12-channel head coil. A structural volume was acquired in the sagittal plane using a magnetization prepared rapid gradient echo (MPRAGE) sequence: TR = 2300 ms, TE = 2.98 ms, FA = 9°, 176 slices, FoV = 256 × 256 mm2, voxel size = 1 × 1 × 1 mm^3^. For functional acquisitions, an echo-planar imaging (EPI) gradient echo sequence was used with the following parameters: TR = 2160 ms, TE = 30 ms, FA = 90°, FoV = 220 × 220 mm^2^, matrix size = 64 × 64, 40 transverse slices, slice thickness = 3 mm, 10% inter-slice gap, inplane resolution = 3.44 × 3.44 mm^2^. In order to minimize the effects of gradient artifact on electroencephalography recordings, the sequence parameters were chosen so that the MR scan repetition time (2160 ms) matched a common multiple of the EEG sample time (0.2 ms), the product of the scanner clock precision (0.1 ms) and the number of slices (40 slices).

EEG was recorded simultaneously with MR using a 64-channel MR-compatible cap (Braincap MR, Easycap, Herrsching, Germany) and two MR-compatible 32-channel amplifiers (Brainamp MR plus, Brain Products GmbH, Gilching, Germany). EEG caps included 62 scalp electrodes referenced to FCz. Two bipolar ECG derivations, V2–V5 and V3 –V6, were recorded using a 16-channel MR-compatible amplifier (Brainamp ExG MR, Brain Products GmbH, Gilching, Germany). Electrode impedances were maintained below 5 kΩ. Data were sampled at 5 kHz with a resolution of 500 nV and analog-filtered between 0.0159 and 250 Hz. Further details of simultaneous EEG–fMRI acquisition have been reported previously ^98^.

#### fMRI datasets: Anesthesia Dataset-SHH

This dataset served as the basis for published studies^15,99–101^. A total of 24 participants (12 male, 46.33 ± 11.39 years) were included in this study (Table 1) and received intravenous propofol anesthesia. All participants received the elective transsphenoidal approach for resection of pituitary microadenoma. Participants were classified as American Society of Anesthesiologists physical status I or II and had no history of craniotomy, cerebral neuropathy, vital organ dysfunction, or contraindication for a MRI examination. Two resting state fMRI scans (8-min each) were acquired in wakefulness and anesthesia state for each participant. Under awake state, participants were instructed to wear earplugs, take a comfortable supine position, relax, eyes closed during scanning, and not to concentrate on anything in particular. The subjects were anesthetized after the awake state scan. Fifteen minutes after stabilization of anesthetic levels and hemodynamics, the fMRI scan in anesthetized state was done. Written informed consent was obtained from all participants, and the study was approved by the Ethics Committee of Shanghai Huashan Hospital, Fudan University, Shanghai, China.

For anesthesia induction, a target-controlled infusion (TCI) was used to set plasma concentration of propofol at 3.0–5.0 μg/ml, followed by remifentanil (1.0 μg/kg) and succinylcholine (1.5 mg/kg) to facilitate endotracheal intubation. The TCI propofol was maintained at a stable effect-site concentration (4.0 μg/ml) to reliably induce an unconscious state. As there is a quick elimination of the analgesic remifentanil and the depolarized neuromuscular relaxant succinylcholine from plasma, the anesthetic effects on the brain are considered to be solely pertaining to propofol. After the induction of anesthesia, the subjects were ventilated with intermittent positive pressure ventilation, setting the tidal volume at 8–10 ml/kg, the respiratory rate at 10–12 beats per minute, and maintaining PetCO2 (end tidal partial pressure of CO2) at 35–40 mm Hg. All subjects fulfilled the criteria of deep sedation, and showed neither response to verbal command (“strongly squeeze my hand!”) nor response to prodding or shaking during anesthesia, corresponding to Ramsay 5–6^102^ and an OAA/S score of 1^103^. No subject reported explicit memory in the post-operative follow-up. Therefore, all subjects were considered unconscious during anesthesia. Additional details are provided in the previous report^100,101,104^.

A Siemens 3 T scanner (Siemens, Erlangen, Germany) was used to obtain functional images of the whole brain using a T2*-weighted EPI series (TR/TE/θ = 2000 ms/30 ms/90◦, FOV = 192 mm, matrix size = 64 × 64, gap = 0 mm). A total of 240 volumes were acquired for each resting-state scan. A high resolution T1-weighted anatomical image was acquired for each participant for functional image registration and localization.

#### fMRI datasets: UWS Dataset-ZJH

This dataset served as the basis for published studies^5,99,105^. We included 20 UWS patients (11 male, age 51.60 ± 10.78 years) with relatively well-preserved brain structures and 20 healthy subjects (10 male, age 32.35 ± 10.69) (Table 1). The UWS patients were assessed using the Coma Recovery Scale-Revised (CRS-R) ^106^. Healthy participants reported no history of neurological or psychiatric disorders. Resting-state fMRI scan (∼8 min) was acquired for each participant. During scanning, participants were instructed to lie supine, remain relaxed, and not focus on anything in particular. Informed written consent was obtained from all healthy participants and the legal representatives of the patients. The study was approved by the Ethics Committee of Zhujiang Hospital, Guangzhou, China.

Image data were acquired on the same Philips 3 Tesla scanner. Functional images were acquired using a T2*-weighted EPI sequence: TR/TE = 2000 ms/30 ms, FOV = 224 × 224 mm2, matrix = 64 × 64, 33 slices, thickness = 3.5 mm, gap = 0.7 mm. A total of 240 fMRI volumes were acquired. A high-resolution T1-weighted anatomical image was also acquired for all participants with parameters: TE/TR = 3.2/7 ms, slice thickness = 1 mm, matrix size = 256 × 256.

#### EEG datasets: Sleep Dataset-ANPHY

The ANPHY-Sleep dataset is an open-access collection of overnight polysomnographic recordings from 29 healthy adults (13 female, aged 32.17 ± 6.30 years) recruited at the Montreal Neurological Institute^107^. The dataset is publicly available through the Open Science Framework (https://doi.org/10.17605/OSF.IO/R26FH). It contains 83-channel high-density EEG (HD-EEG), ECG, EMG, EOG, manually scored sleep-stage annotations, and average co-registered electrode positions. Participants spent approximately 7 h in the sleep laboratory from 21:00 – 23:00 until 05:00 – 07:00 the next morning. The recordings were scored according to AASM by a certified sleep technician. For the N3 EEG awakening analysis, 60-s N3 segments were categorized according to the subsequent sleep state as N3F-W or N3F-N2. Nineteen participants contributed N3F-W segments and 26 contributed N3F-N2 segments (Table 1). Details of segment selection and spectral analyses are described below.

Overnight PSG and HD-EEG signals were acquired using a JE-120 256-channel EEG/PSG system (Nihon Kohden, Tokyo, Japan). 83 electrodes were positioned according to the international 10-10 system and secured with collodion. Electrode impedances were maintained below 5 kΩ. Four periocular electrodes provided vertical and horizontal EOG recordings, and six electrodes were used for bipolar EMG recordings. A single modified lead II ECG channel was recorded on the torso. All continuous signals were sampled at 1,000 Hz. Further details of the dataset have been reported previously^107^.

#### EEG datasets: UWS Dataset-GWIRH

This dataset was originally collected for a previous study^108^. The original cohort included 16 patients diagnosed with UWS (9 male, mean age: 51.68 ± 11.05 years). Clinical diagnoses were established using the Coma Recovery Scale-Revised (CRS-R). Assessments were conducted one week before the experiment and repeated on the day of data collection. During each assessment period, two experienced physicians performed at least two CRS-R evaluations. For the present analysis, 10 patients (7 male, mean age: 53.10 ± 13.10 years) had both usable N3 EEG data and CRS-R scores at the three-month follow-up and were included in the analysis (Table 1). Among these 10 patients, etiologies included cerebrovascular disease (CVD; n = 4), traumatic brain injury (TBI; n = 4), and anoxic brain injury (ABI; n = 2). Ethical approval was obtained and informed consent was acquired from legal guardians.

Overnight polysomnography was acquired using a Compumedics Grael PSG system with Profusion software (version 4.5, build 500; Compumedics Limited). The mean recording duration was 8.39 ± 0.41 h. Recordings included EEG, EOG, EMG, and synchronized video. EEG was recorded from six electrodes positioned according to the international 10–20 system (F3, F4, C3, C4, O1, and O2) at 256 Hz. Electrode impedances were maintained below 5 k Ω. Synchronized video was used to annotate eye opening, rest, agitation, and nursing-related movements.

Given the abnormal sleep patterns in UWS patients, the standard sleep-scoring criteria had to be adapted to these patients’ sleep patterns^109–111^. All sleep data were independently scored by two expert clinical neurophysiologists. Discrepancies in sleep scoring and arousal recognition were reviewed and resolved through consensus. Uncertain epochs were excluded from analyses to maintain data integrity. Further details of the dataset and criteria of sleep scoring have been reported previously^112^.

#### Image preprocessing

Functional MRI data from simultaneous EEG-fMRI and fMRI datasets were preprocessed using afni_proc.py in AFNI (https://afni.nimh.nih.gov/) with anatomical segmentations derived from FreeSurfer (https://surfer.nmr.mgh.harvard.edu/fswiki). Preprocessing included removal of the first three volumes, elimination of spikes, slice-timing correction, EPI–anatomical alignment, motion correction, spatial smoothing, scaling, and nuisance regression. Functional images were aligned to individual anatomical image without normalization to a standard space, and registered to the volume with the minimum outlier fraction, then resampled to 3 mm isotropic voxels and smoothed with a 4 mm FWHM Gaussian kernel. FreeSurfer-derived ventricular and white matter masks were used for tissue-based denoising. Nuisance regression included demeaned motion parameters, motion derivatives, the first five principal components extracted from the ventricular signal, ANATICOR-based local white matter regression, and temporal band-pass filtering (0.01–0.1 Hz) implemented within the regression model. Volumes with framewise displacement > 0.4 mm, as well as preceding volume, and volumes with >10% outlier voxels were replaced by temporal interpolation.

#### ROI definition

Regions of interest (ROIs) were defined in each participant’s native anatomical space and mapped to the preprocessed functional space. Cortical ROIs were based on the Schaefer2018 parcellation (100 parcels, 7 networks)^113^, which was selected because its relatively coarse resolution is better suited to UWS patients with structural distortion and facilitates extraction of signals from preserved cortical regions. The parcellation was projected from template surface space to individual subject space and converted to volumetric space. Thalamic ROIs were derived from subject-specific thalamic nuclei segmentations generated using the probabilistic atlas implemented in FreeSurfer, which provides voxel-wise segmentation probabilities in native anatomical space^114^. The four pulvinar subregions were combined into a single pulvinar nucleus (PUL), and the two mediodorsal subregions were combined into a single mediodorsal nucleus (MD)^21^. In total, 18 thalamic ROIs were included bilaterally: anteroventral nucleus (AV), CM, lateral geniculate nucleus (LGN), MD, PUL, ventral anterior nucleus (VA), ventral lateral anterior nucleus (VLA), ventral lateral posterior nucleus (VLP), and VPL. Mean time series were extracted from each cortical and thalamic ROI, yielding 118 ROIs in total for subsequent analyses.

#### Edge-centric analysis: Edge community construction

Edge-centric analysis was performed as described previously^56–58^. For each participant, edge time series (ETS) were constructed by first z-scoring the time series of all nodes and then computing the element-wise product for every pair of nodes (*i* and *j, i*≠*j*). The resultant ETS matrix 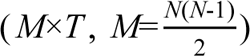 encodes the moment-by-moment co-fluctuations of edges. ETS matrix was then clustered using k-means algorithm (250 replicates; maximum 1,000 iterations), assigning community labels for all edges. The resulting edge-community assignments were mapped back onto an *N*×*N* symmetric matrix. To assess robustness across clustering resolutions, the number of clusters (*K*) was varied from 3 to 20 in increment of 1. The main results are reported for = 10, consistent with prior studies^56,67,68^.

#### Edge-centric analysis: Community similarity

Community similarity was used to quantify the similarity of edge-community profiles between pairs of nodes. In the edge-community matrix *X* (*N*×*N*), *x_ij_* denotes the community assignment of the edge between nodes *i* and *j*. Accordingly, each row (or column) of *X* represented the edge-community profile of one node. Community similarity between nodes i and j was defined as the proportion of the same community labels across their edge profiles as below:

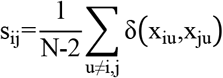

Where δ(x_iu_,x_ju_) is the Kronecker delta, which equals 1 when x_iu_=x_ju_ and 0 otherwise. Higher community similarity (s_ij_) indicates that nodes *i* and *j* exhibit more similar edge-community participation, whereas lower value indicates more distinct edge-community profiles. Estimating community similarity between all node pairs yielded an *N*×*N* similarity matrix (118 × 118 in this study, Fig. 1).

For each thalamic nucleus, thalamic community similarity was computed by averaging its community similarity with all other thalamic nuclei after excluding self-similarity. Similarity from left and right homologous nuclei were then averaged to obtain bilateral nucleus-wise community similarity. For each cortical network, within-network community similarity was calculated by averaging all pairwise community similarity among ROIs belonging to that network.

#### Edge-centric analysis: Normalized entropy

Normalized entropy was used to quantify the diversity of edge-community participation for each node. After assigning all edges to *k* communities, the participation of node *i* in community *c* was defined as

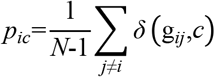

Where *N* is the total number of nodes, g_ij_∈{1,…,k} denotes the community assignment of the edge between nodes i and j, and δ(g_ij,c_) is the Kronecker delta, which equals 1 when g_ij_=c and 0 otherwise. Thus, p_ic_ represents the proportion of edges connected to node *i* that belong to community c. The entropy of the edge-community participation distribution for node *i, p_i_* =[*p_i_*_1_, …*p_ik_*], was then computed as

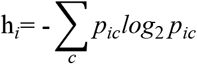

To facilitate comparison across different numbers of communities, entropy was normalized by *log*_2_*k*, thereby constraining the metric between 0 and 1. Higher normalized entropy indicates that the edges of a node are distributed more evenly across multiple communities, whereas lower value indicates that the edges are concentrated within fewer communities.

For thalamic nuclei, normalized entropy was first obtained for each of the 18 unilateral thalamic ROIs, after which values from left and right homologous nuclei were averaged to obtain bilateral nucleus-wise normalized entropy. For cortical networks, within-network averaged entropy was calculated by averaging normalized entropy across all ROIs belonging to each canonical network.

#### Edge-centric analysis: Triad motifs

We used graph-based motif analysis to quantify higher-order community organization from the edge-community assignments. A triad motif was defined by a reference node and two additional nodes, with motif type determined by the community labels of the three constituent edges (Fig. 3A). Four motif types were considered: closed loop, in which all three edges share the same label; forked, in which the two edges connected to the reference node share the same label but the third differs; L-shape, in which one reference-linked edge shares its label with the third edge whereas the other differs; and diverse, in which all three edges belong to different communities. For each reference node, we calculated the fraction of triads assigned to each motif type.

In the thalamic–thalamic analysis, the reference node was a thalamic nucleus and the other two nodes were selected from the remaining thalamic nuclei. In the thalamic–cortical analysis, the reference node was a thalamic nucleus and the other two nodes were selected from cortical regions. Fractions from left and right homologous nuclei were averaged to obtain bilateral nucleus-wise motif fractions.

#### NPI inferred effective connectivity

NPI is a data-driven framework that estimates causal interactions by virtually perturbing a surrogate model of brain dynamics and was used here to infer directed effective connectivity from fMRI ROI signals^59^. For each participant, an artificial neural network was trained as a surrogate model of large-scale brain dynamics, such that the neural state at time t+1 was predicted from the preceding three time points:

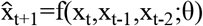

where x_t_ denotes the neural state at time t, θ represents the parameters of the multilayer perceptron (MLP) implemented as the artificial neural network (function *f*), and *x^_t_*_+1_ denotes the predicted state at time *t*+1.

After model training, directed effective connectivity from source node *i* to all other nodes was estimated by applying a virtual perturbation on the node *i* and quantifying the changes in predicted activity across all nodes relative to the unperturbed baseline:

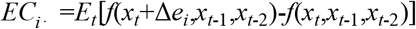

where *e_i_* is a unit vector indicating the perturbed node, Δ is the perturbation magnitude, and *EC_i_*_⋅_ represents effective connectivity from node *i* to all other regions. Averaging perturbation responses across time points yielded a participant-level directed effective connectivity matrix, and group-level effective connectivity was obtained by averaging participant-level matrices. The resulting *EC* matrix is of size *N*×*N*, where the *i*-th row represents the effective connectivity from region *i* to all other regions, and the *j*-th column represents the effective connectivity from all other regions to region *j* (Fig. 1). Positive and negative values indicate excitatory and inhibitory effective connectivity, respectively.

To minimize dependence on any single perturbation magnitude, effective connectivity was averaged across perturbation strengths from 1 to 10 (step size = 1). Effective connectivity of left and right homologous nuclei were subsequently averaged to derive bilateral nucleus-wise effective connectivity.

To evaluate how well the surrogate models reproduced empirical functional connectivity (FC), we calculated empirical FC from the observed BOLD signals and model-derived FC from the trained surrogate models for each participant. The resulting FC matrices were averaged across participants within each state, and Pearson correlations were then calculated between the non-diagonal elements of the group-averaged empirical and model-derived FC matrices.

#### Statistical analysis of fMRI data

Linear mixed-effects models (LMMs) were used for Sleep Dataset-WU and Sleep Dataset-IUGM to assess differences across wakefulness, N2 and N3 sleep. In these models, sleep stage was included as a fixed effect and subject as a random intercept. Post hoc pairwise comparisons were further performed on estimated marginal means using two-sided Wald-z tests.

For Anesthesia Dataset-SHH, comparisons between wakefulness and propofol-induced deep sedation were performed using two-sided paired-sample tests because both conditions were acquired from the same participants. Normality of the paired differences was assessed using the Shapiro-Wilk test. Paired-sample t-tests were used when the paired differences were normally distributed, whereas Wilcoxon signed-rank tests were used when the paired differences were non-normally distributed. For UWS Dataset-ZJH, comparisons between healthy controls and patients with UWS were performed using two-sided independent-sample tests. Normality was assessed using the Shapiro-Wilk test and homogeneity of variance using Levene’s test. Student’s t-tests were used when the assumptions of normality and equal variance were met, Welch’s t-tests when variances were unequal, and Mann– Whitney U tests when the normality assumption was not met.

Bonferroni correction was applied separately within each predefined analysis family, with statistical significance set at ɑ = 0.05. For whole-thalamus analyses of community similarity and normalized entropy, correction was performed across nine thalamic nuclei. For cortical community similarity and normalized entropy, correction was performed across seven cortical networks. For CM- and VPL-centered triadic motif analyses, correction was performed across the four motif types within each reference nucleus. For NPI analyses, correction was performed across the two predefined directed connections, CM→VPL and VPL→PUL. For post hoc comparisons following LMMs, p values were corrected across the three pairwise sleep-stage contrasts (W vs N2, W vs N3 and N2 vs N3) within each feature. For the independent Sleep Dataset-IUGM, the LMM analysis of community similarity was corrected across the two prespecified target nuclei, CM and VPL, whereas all other correction procedures were identical to those used for Sleep Dataset-WU.

#### EEG preprocessing

EEG data from simultaneous EEG-fMRI datasets (Sleep Dataset-WU and Sleep Dataset-IUGM) were preprocessed according to their respective published pipelines^97,98^. Both pipelines included adaptive average subtraction of MR gradient artifacts, downsampling to 250 Hz, 60-Hz low-pass filtering, and rereferencing to averaged mastoids. Ballistocardiographic (BCG) artifacts time-locked to ECG R peaks were attenuated using template subtraction and independent component analysis (ICA). Gradient-artifact correction was performed using Brain Products Analyzer for Dataset-WU and the FASST toolbox for Dataset-IUGM. In Dataset-WU, R peaks were visually verified and manually corrected when necessary, with additional ICA applied to residual BCG artifacts. In Dataset-IUGM, BCG correction used an algorithm combining template subtraction with event-related ICA. Further preprocessing details have been reported previously^97,98^.

EEG data from Sleep Dataset-ANPHY were high-pass filtered at 0.3 Hz, low-pass filtered at 45 Hz, and downsampled from 1,000 to 250 Hz. The signals were then rereferenced to the average of the bilateral mastoid electrodes (TP9 and TP10).

EEG data from UWS Dataset-GWIRH were notch filtered at 50 Hz and low-pass filtered at 45 Hz. The signals were rereferenced to the global average of all retained channels. Expert clinical neurophysiologists visually inspected the data and manually excluded contaminated segments, using an amplitude threshold of ±100 μV as an artifact-rejection criterion. Each channel was z-score normalized to reduce interchannel variability. Finally, the signals were downsampled to 100 Hz.

#### Association between EEG delta power and fMRI effective connectivity during N3 sleep

To assess the association between EEG delta power and fMRI effective connectivity during N3 sleep, we extracted EEG segments corresponding temporally to each fMRI N3 segment included in the analysis. For each retained EEG channel, power spectral density was estimated using Welch’s method with 2-s Hamming windows and 50% overlap. Relative delta power was calculated as delta-band power (1 – 4 Hz) divided by total spectral power (1 – 40 Hz) and then averaged across channels. Data from Sleep Dataset-WU and Sleep Dataset-IUGM were pooled. Partial Pearson correlations assessed the associations of relative delta power with fMRI-derived effective connectivity from CM to VPL and from VPL to PUL, with dataset identity included as a covariate. The two tests were corrected for multiple comparisons using the Bonferroni method.

#### Comparison of relative delta power in N3 segments followed by wakefulness and N2 sleep

Relative delta power was quantified over the final 60s of each N3 segment, classified according to whether the next scored sleep stage was wakefulness (N3F-W) or N2 sleep (N3F-N2). A fixed 60-s window was used to ensure comparable analysis durations across datasets, and segments without a complete 60-s window were excluded. For each retained EEG channel, power spectral density was estimated using Welch’s method with 2-s Hamming windows and 50% overlap. Relative delta power was calculated as spectral power within 1-4 Hz divided by total power within 1-40 Hz and was subsequently averaged across retained EEG channels. Sleep Dataset-WU was excluded because it contained only one N3F-W segment; the analysis was therefore restricted to Sleep Dataset-IUGM (n_N3F-W_ = 7, n_N3F-N2_ = 17) and Sleep Dataset-ANPHY (n_N3F-W_ = 44, n_N3F-N2_ = 148).

Within each dataset, segment-level relative delta power was compared between N3F-W and N3F-N2 using LMM. Subsequent sleep stage was included as a fixed effect, and participant was included as a random intercept to account for multiple segments contributed by the same participant. A one-tailed test was used because we hypothesized that relative delta power would be higher in N3F-W than in N3F-N2.

#### Correlation between relative delta power in N3 and CRS-R change in UWS patients

In UWS Dataset-GWIRH, preprocessed EEG data during N3 were extracted for subsequent analysis. For each retained EEG channel, power spectral density was calculated using Welch’s method with 2-s Hamming windows and 50% overlap. Relative delta power was defined as spectral power within 1-4 Hz divided by total power within 1-40 Hz and was subsequently averaged across retained EEG channels, yielding one N3 relative delta power estimate per UWS patient.

N3 relative delta power was then correlated with clinical data. Clinical improvement was quantified as the change in CRS-R score from baseline to the three-month follow-up, calculated as the follow-up score minus the baseline score. Partial spearman correlation between N3 relative delta power and CRS-R change was calculated while controlling for injury etiology. Statistical significance was evaluated using two-tailed test.

