## SupplementaryMaterial for "Intra-thalamic coordination maintains arousability in slow-wave sleep"

**Table S1. Effect of sleep stages on thalamic community similarity in Sleep Dataset-WU**

| ROI | F | DF1 | DF2 | p | $p_{Bonf}$ |
|---|---|---|---|---|---|
| AV | 0.043 | 2.000 | 31.233 | 0.958 | 1.000 |
| CM | 6.791 | 2.000 | 51.000 | 0.002 | 0.022* |
| LGN | 1.857 | 2.000 | 34.944 | 0.171 | 1.000 |
| MD | 4.373 | 2.000 | 30.949 | 0.021 | 0.191 |
| PUL | 4.940 | 2.000 | 36.680 | 0.013 | 0.113 |
| VA | 0.046 | 2.000 | 31.903 | 0.955 | 1.000 |
| VLA | 3.606 | 2.000 | 32.304 | 0.039 | 0.347 |
| VLP | 5.024 | 2.000 | 32.624 | 0.013 | 0.113 |
| VPL | 6.379 | 2.000 | 34.382 | 0.004 | 0.040* |

Asterisks indicate significance after Bonferroni correction

**Table S2. Post hoc comparisons for thalamic community similarity with significant sleep-stage effects in Sleep Dataset-WU**

| ROI | Level 1 | Level 2 | Estimate | SE | DF | z | p | $p_{Bonf}$ |
|---|---|---|---|---|---|---|---|---|
| CM | W | N2 | -0.180 | 0.056 | Inf | -3.199 | 0.001 | 0.004** |
| CM | W | N3 | -0.221 | 0.072 | Inf | -3.057 | 0.002 | 0.007** |
| CM | N2 | N3 | -0.041 | 0.069 | Inf | -0.589 | 0.556 | 1.000 |
| VPL | W | N2 | -0.155 | 0.050 | Inf | -3.133 | 0.002 | 0.005** |
| VPL | W | N3 | -0.194 | 0.066 | Inf | -2.923 | 0.004 | 0.010* |
| VPL | N2 | N3 | -0.038 | 0.063 | Inf | -0.610 | 0.542 | 1.000 |

Asterisks indicate significance after Bonferroni correction; all tests were two-sided.

**Table S3. Comparisons of thalamic community similarity between wakefulness and deep sedation in Anesthesia Dataset-SHH**

| ROI | p for Shapiro-Wilk normality test | statistic | df | p | $p_{Bonf}$ |
|---|---|---|---|---|---|
| AV | 0.097 | 2.030[a] | 23 | 0.054 | 0.487 |
| CM | 0.114 | 0.754[a] | 23 | 0.459 | 1.000 |
| LGN | 0.049 | 168.500[b] | - | 0.608 | 1.000 |
| MD | 0.097 | 1.798[a] | 23 | 0.085 | 0.767 |
| PUL | 0.067 | 1.609[a] | 23 | 0.121 | 1.000 |
| VA | 0.101 | 2.286[a] | 23 | 0.032 | 0.286 |
| VLA | 0.023 | 250.000[b] | - | 0.003 | 0.028* |
| VLP | 0.006 | 248.000[b] | - | 0.004 | 0.035* |
| VPL | 0.031 | 195.000[b] | - | 0.208 | 1.000 |

a = t statistic from Student’s t-test; b = W statistic from Wilcoxon signed-rank test; asterisks indicate significance after Bonferroni correction; all tests were two-sided.

**Table S4. Comparisons of thalamic community similarity between HC and UWS in UWS Dataset-ZJH**

| ROI | p for Shapiro-Wilk normality test | | p for Levene's test for equal variances | statistic | df | p | $p_{Bonf}$ |
|---|---|---|---|---|---|---|---|
| | HC | UWS | | | | | |
| AV | 0.099 | 0.996 | 0.430 | 3.346[a] | 38 | 0.002 | 0.017* |
| CM | 0.132 | 0.049 | 0.764 | 477.000[b] | - | 0.072 | 0.646 |
| LGN | 0.273 | 0.175 | 0.494 | 0.100[a] | 38 | 0.921 | 1.000 |
| MD | 0.192 | 0.757 | 0.148 | 5.192[a] | 38 | < 0.001 | < 0.001*** |
| PUL | 0.630 | 0.672 | 0.477 | 2.530[a] | 38 | 0.016 | 0.141 |
| VA | 0.140 | 0.421 | 0.662 | 4.680[a] | 38 | < 0.001 | < 0.001*** |
| VLA | 0.059 | 0.335 | 0.026 | 4.054[c] | 33.737 | < 0.001 | 0.003** |
| VLP | 0.177 | 0.235 | 0.027 | 3.754[c] | 34.317 | 0.001 | 0.006** |
| VPL | 0.270 | 0.566 | 0.715 | 2.376[a] | 38 | 0.023 | 0.204 |

a = t statistic from Student's t-test; b = W statistic from Mann–Whitney U test; c = t statistic from Welch's t-test; asterisks indicate significance after Bonferroni correction; all tests were two-sided.

**Table S5. Effect of sleep stages on network community similarity in Sleep Dataset-WU**

| ROI | F | DF1 | DF2 | p | $p_{Bonf}$ |
|---|---|---|---|---|---|
| VN | 2.389 | 2.000 | 37.821 | 0.105 | 0.738 |
| SMN | 3.182 | 2.000 | 31.711 | 0.055 | 0.385 |
| DAN | 9.517 | 2.000 | 32.462 | 0.001 | 0.004** |
| VAN | 3.807 | 2.000 | 28.899 | 0.034 | 0.239 |
| LN | 5.158 | 2.000 | 51.000 | 0.009 | 0.064 |
| FPN | 0.129 | 2.000 | 35.792 | 0.880 | 1.000 |
| DMN | 1.464 | 2.000 | 36.762 | 0.245 | 1.000 |

Asterisks indicate significance after Bonferroni correction.

**Table S6. Post hoc comparisons for network community similarity with significant sleep-stage effects in Sleep Dataset-WU**

| ROI | Level 1 | Level 2 | Estimate | SE | DF | z | p | $p_{Bonf}$ |
|---|---|---|---|---|---|---|---|---|
| DAN | W | N2 | 0.086 | 0.024 | Inf | 3.574 | < 0.001 | 0.001** |
| DAN | W | N3 | 0.123 | 0.032 | Inf | 3.824 | < 0.001 | < 0.001*** |
| DAN | N2 | N3 | 0.037 | 0.030 | Inf | 1.215 | 0.225 | 0.674 |

Asterisks indicate significance after Bonferroni correction; all tests were two-sided.

**Table S7. Comparisons of network community similarity between wakefulness and deep sedation in Anesthesia Dataset-SHH**

| ROI | p for Shapiro-Wilk normality test | statistic | df | p | $p_{Bonf}$ |
|---|---|---|---|---|---|
| VN | < 0.001 | 166.5[b] | - | 0.649 | 1.000 |
| SMN | 0.376 | 3.549[a] | 23 | 0.002 | 0.012* |
| DAN | 0.324 | 5.181[a] | 23 | < 0.001 | < 0.001*** |
| VAN | 0.391 | 2.680[a] | 23 | 0.013 | 0.094 |
| LN | 0.740 | 0.337[a] | 23 | 0.739 | 1.000 |
| FPN | 0.797 | -0.018[a] | 23 | 0.986 | 1.000 |
| DMN | 0.913 | -0.98[a] | 23 | 0.337 | 1.000 |

a = t statistic from Student's t-test; b = W statistic from Wilcoxon signed-rank test; asterisks indicate significance after Bonferroni correction; all tests were two-sided.

**Table S8. Comparisons of network community similarity between HC and UWS in UWS Dataset-ZJH**

| ROI | p for Shapiro-Wilk normality test | | p for Levene's test for equal variances | statistic | df | p | $p_{Bonf}$ |
|---|---|---|---|---|---|---|---|
| | HC | UWS | | | | | |
| VN | 0.252 | 0.017 | 0.496 | 520.000[b] | - | 0.002 | 0.017* |
| SMN | 0.312 | 0.339 | 0.029 | 5.915[c] | 33.463 | < 0.001 | < 0.001*** |
| DAN | 0.893 | 0.101 | 0.268 | 3.245[a] | 38.000 | 0.003 | 0.017* |
| VAN | 0.991 | 0.026 | 0.607 | 578.500[b] | - | < 0.001 | < 0.001*** |
| LN | 0.002 | 0.954 | 0.858 | 507.000[b] | - | 0.008 | 0.056 |
| FPN | 0.002 | 0.034 | 0.020 | 415.000[b] | - | 0.904 | 1.000 |
| DMN | 0.001 | 0.030 | 0.756 | 527.000[b] | - | 0.001 | 0.008** |

a = t statistic from Student's t-test; b = W statistic from Mann–Whitney U test; c = t statistic from Welch's t-test; asterisks indicate significance after Bonferroni correction; all tests were two-sided.

**Table S9. Effect of sleep stages on thalamic normalized entropy in Sleep Dataset-WU**

| ROI | F | DF1 | DF2 | p | $p_{Bonf}$ |
|---|---|---|---|---|---|
| AV | 2.592 | 2.000 | 39.108 | 0.088 | 0.789 |
| CM | 1.638 | 2.000 | 33.981 | 0.209 | 1.000 |
| LGN | 1.326 | 2.000 | 33.590 | 0.279 | 1.000 |
| MD | 0.880 | 2.000 | 38.673 | 0.423 | 1.000 |
| PUL | 0.185 | 2.000 | 51.000 | 0.832 | 1.000 |
| VA | 0.982 | 2.000 | 51.000 | 0.381 | 1.000 |
| VLA | 2.075 | 2.000 | 51.000 | 0.136 | 1.000 |
| VLP | 0.987 | 2.000 | 39.233 | 0.382 | 1.000 |
| VPL | 0.641 | 2.000 | 34.525 | 0.533 | 1.000 |

Asterisks indicate significance after Bonferroni correction.

**Table S10. Comparisons of thalamic normalized entropy between wakefulness and deep sedation in Anesthesia Dataset-SHH**

| ROI | p for Shapiro-Wilk normality test | statistic | df | p | $p_{Bonf}$ |
|---|---|---|---|---|---|
| AV | 0.157 | -0.212[a] | 23 | 0.834 | 1.000 |
| CM | 0.882 | -0.607[a] | 23 | 0.550 | 1.000 |
| LGN | 0.676 | 1.306[a] | 23 | 0.204 | 1.000 |
| MD | 0.299 | -0.702[a] | 23 | 0.490 | 1.000 |
| PUL | 0.234 | 1.678[a] | 23 | 0.107 | 0.962 |
| VA | 0.586 | 0.518[a] | 23 | 0.609 | 1.000 |
| VLA | 0.809 | 1.028[a] | 23 | 0.315 | 1.000 |
| VLP | 0.387 | 0.783[a] | 23 | 0.442 | 1.000 |
| VPL | 0.834 | 0.525[a] | 23 | 0.605 | 1.000 |

a = t statistic from Student's t-test; all tests were two-sided.

**Table S11. Comparisons of thalamic normalized entropy between HC and UWS in UWS Dataset-ZJH**

| ROI | p for Shapiro-Wilk normality test | | p for Levene's test for equal variances | statistic | df | p | $p_{Bonf}$ |
|---|---|---|---|---|---|---|---|
| | HC | UWS | | | | | |
| AV | 0.552 | 0.088 | 0.736 | -0.836[a] | 38 | 0.409 | 1.000 |
| CM | 0.416 | 0.238 | 0.439 | 0.983[a] | 38 | 0.332 | 1.000 |
| LGN | 0.004 | 0.423 | 0.422 | 456.000[b] | - | 0.221 | 1.000 |
| MD | 0.758 | 0.716 | 0.876 | -0.399[a] | 38 | 0.692 | 1.000 |
| PUL | 0.062 | 0.056 | 0.309 | 0.999[a] | 38 | 0.324 | 1.000 |
| VA | 0.957 | 0.335 | 0.386 | -0.404[a] | 38 | 0.689 | 1.000 |
| VLA | 0.249 | 0.170 | 0.878 | 0.061[a] | 38 | 0.952 | 1.000 |
| VLP | 0.035 | 0.148 | 0.897 | 422.000[b] | - | 0.758 | 1.000 |
| VPL | 0.076 | 0.345 | 0.855 | 0.660[a] | 38 | 0.514 | 1.000 |

a = t statistic from Student's t-test; b = W statistic from Mann-Whitney U test; c = t statistic from Welch's t-test; asterisks indicate significance after Bonferroni correction; all tests were two-sided.

**Table S12. Effect of sleep stages on network normalized entropy in Sleep Dataset-WU**

| ROI | F | DF1 | DF2 | p | $p_{Bonf}$ |
|---|---|---|---|---|---|
| VN | 4.173 | 2.000 | 37.759 | 0.023 | 0.161 |
| SMN | 1.612 | 2.000 | 33.451 | 0.215 | 1.000 |
| DAN | 14.459 | 2.000 | 33.924 | < 0.001 | < 0.001*** |
| VAN | 0.769 | 2.000 | 35.376 | 0.471 | 1.000 |

| | | | | | |
|---|---|---|---|---|---|
| LN | 1.187 | 2.000 | 38.162 | 0.316 | 1.000 |
| FPN | 4.157 | 2.000 | 51.000 | 0.021 | 0.149 |
| DMN | 3.593 | 2.000 | 37.913 | 0.037 | 0.261 |

Asterisks indicate significance after Bonferroni correction.

**Table S13. Post hoc comparisons for networks with significant sleep-stage effects in Sleep Dataset-WU**

| ROI | Level 1 | Level 2 | Estimate | SE | DF | z | p | $p_{Bonf}$ |
|---|---|---|---|---|---|---|---|---|
| DAN | W | N2 | 0.073 | 0.014 | Inf | 5.173 | < 0.001 | < 0.001*** |
| DAN | W | N3 | 0.070 | 0.019 | Inf | 3.603 | < 0.001 | 0.001** |
| DAN | N2 | N3 | -0.004 | 0.018 | Inf | -0.213 | 0.831 | 1.000 |

Asterisks indicate significance after Bonferroni correction; all tests were two-sided.

**Table S14. Comparisons of network normalized entropy between wakefulness and deep sedation in Anesthesia Dataset-SHH**

| ROI | p for Shapiro-Wilk normality test | statistic | df | p | pBonf |
|---|---|---|---|---|---|
| VN | 0.552 | $-1.048^{a}$ | 23 | 0.306 | 1.000 |
| SMN | 0.341 | $2.106^{a}$ | 23 | 0.046 | 0.325 |
| DAN | 0.852 | $0.661^{a}$ | 23 | 0.515 | 1.000 |
| VAN | 0.952 | $-1.885^{a}$ | 23 | 0.072 | 0.505 |
| LN | 0.259 | $-1.889^{a}$ | 23 | 0.072 | 0.501 |
| FPN | 0.051 | $-3.723^{a}$ | 23 | 0.001 | 0.008** |
| DMN | 0.768 | $-5.715^{a}$ | 23 | < 0.001 | < 0.001*** |

a = t statistic from Student's t-test; asterisks indicate significance after Bonferroni correction; all tests were two-sided.

**Table S15. Comparisons of network normalized entropy between HC and UWS in UWS Dataset-ZJH**

| ROI | p for Shapiro-Wilk normality test | | p for Levene's test for equal variances | statistic | df | p | pBonf |
|---|---|---|---|---|---|---|---|
| | HC | UWS | | | | | |
| VN | 0.055 | 0.019 | 0.975 | $320.000^{b}$ | - | 0.014 | 0.100 |
| SMN | 0.498 | 0.910 | 0.679 | $1.616^{a}$ | 38.000 | 0.114 | 0.800 |
| DAN | 0.394 | 0.459 | 0.073 | $-1.719^{a}$ | 38.000 | 0.094 | 0.657 |
| VAN | 0.832 | 0.445 | 0.077 | $1.100^{a}$ | 38.000 | 0.278 | 1.000 |
| LN | 0.461 | 0.271 | 0.414 | $-3.187^{a}$ | 38.000 | 0.003 | 0.020* |
| FPN | 0.540 | 0.057 | 0.486 | $-6.784^{a}$ | 38.000 | < 0.001 | < 0.001*** |
| DMN | 0.911 | 0.943 | 0.040 | $-5.479^{c}$ | 31.039 | < 0.001 | < 0.001*** |

a = t statistic from Student's t-test; b = W statistic from Mann-Whitney U test; c = t statistic from Welch's t-test; asterisks indicate significance after Bonferroni correction; all tests were two-sided.

**Table S16. Effect of sleep stages on the fraction of thalamic-thalamic motif types in Sleep Dataset-WU**

| Motif | F | DF1 | DF2 | p | $p_{Bonf}$ |
|---|---|---|---|---|---|
| **CM and other thalamic nuclei** | | | | | |
| CM-Closed loop | 5.529 | 2.000 | 51.000 | 0.007 | 0.027* |
| CM-Forked | 5.129 | 2.000 | 51.000 | 0.009 | 0.037* |
| CM-L-shape | 1.174 | 2.000 | 33.523 | 0.322 | 1.000 |
| CM-Diverse | 3.685 | 2.000 | 51.000 | 0.032 | 0.128 |
| **VPL and other thalamic nuclei** | | | | | |
| VPL-Closed loop | 5.173 | 2.000 | 36.168 | 0.011 | 0.042* |
| VPL-Forked | 1.443 | 2.000 | 34.115 | 0.250 | 1.000 |
| VPL-L-shape | 3.100 | 2.000 | 38.035 | 0.057 | 0.226 |
| VPL-Diverse | 3.061 | 2.000 | 37.353 | 0.059 | 0.235 |

Asterisks indicate significance after Bonferroni correction.

**Table S17. Post hoc comparisons for thalamic-thalamic motif types showing significant sleep-stage effects in Sleep Dataset-WU**

| Motif | Level 1 | Level 2 | Estimate | SE | DF | z | p | $p_{Bonf}$ |
|---|---|---|---|---|---|---|---|---|
| **CM and other thalamic nuclei** | | | | | | | | |
| CM-Closed loop | W | N2 | -0.235 | 0.082 | Inf | -2.876 | 0.004 | 0.012* |
| CM-Closed loop | W | N3 | -0.291 | 0.105 | Inf | -2.770 | 0.006 | 0.017* |
| CM-Closed loop | N2 | N3 | -0.056 | 0.101 | Inf | -0.553 | 0.581 | 1.000 |
| CM-Forked | W | N2 | 0.113 | 0.039 | Inf | 2.895 | 0.004 | 0.011* |
| CM-Forked | W | N3 | 0.126 | 0.050 | Inf | 2.511 | 0.012 | 0.036* |
| CM-Forked | N2 | N3 | 0.013 | 0.048 | Inf | 0.267 | 0.790 | 1.000 |
| **VPL and other thalamic nuclei** | | | | | | | | |
| VPL-Closed loop | W | N2 | -0.198 | 0.072 | Inf | -2.774 | 0.006 | 0.017* |
| VPL-Closed loop | W | N3 | -0.256 | 0.095 | Inf | -2.687 | 0.007 | 0.022* |
| VPL-Closed loop | N2 | N3 | -0.058 | 0.090 | Inf | -0.639 | 0.523 | 1.000 |

Asterisks indicate significance after Bonferroni correction; all tests were two-sided.

**Table S18. Comparisons of the fraction of thalamic-thalamic motif types between wakefulness and deep sedation in Anesthesia Dataset-SHH**

| Motif | p for Shapiro-Wilk normality test | statistic | df | p | $p_{Bonf}$ |
|---|---|---|---|---|---|
| **CM and other thalamic nuclei** | | | | | |
| CM-Closed loop | 0.063 | 0.915[a] | 23.000 | 0.370 | 1.000 |
| CM-Forked | 0.184 | 1.472[a] | 23.000 | 0.155 | 0.618 |
| CM-L-shape | 0.361 | -0.812[a] | 23.000 | 0.425 | 1.000 |

| | | | | | |
|---|---|---|---|---|---|
| CM-Diverse | 0.525 | -1.969[a] | 23.000 | 0.061 | 0.244 |
| **VPL and other thalamic nuclei** | | | | | |
| VPL-Closed loop | 0.129 | 0.427[a] | 23.000 | 0.674 | 1.000 |
| VPL-Forked | 0.586 | 1.751[a] | 23.000 | 0.093 | 0.373 |
| VPL-L-shape | 0.985 | -1.136[a] | 23.000 | 0.268 | 1.000 |
| VPL-Diverse | 0.629 | -0.972[a] | 23.000 | 0.341 | 1.000 |

a = t statistic from Student's t-test; asterisks indicate significance after Bonferroni correction; all tests were two-sided.

**Table S19. Comparisons of the fraction of thalamic-thalamic motif types between HC and UWS in UWS Dataset-ZJH**

| Motif | p for Shapiro-Wilk normality test | | p for Levene's test for equal variances | statistic | df | p | $p_{Bonf}$ |
|---|---|---|---|---|---|---|---|
| | HC | UWS | | | | | |
| **CM and other thalamic nuclei** | | | | | | | |
| CM-Closed loop | 0.127 | 0.066 | 0.624 | 1.008[a] | 38.000 | 0.320 | 1.000 |
| CM-Forked | 0.039 | 0.123 | 0.066 | 396.000[b] | - | 0.713 | 1.000 |
| CM-L-shape | 0.054 | 0.698 | 0.495 | -0.330[a] | 38.000 | 0.743 | 1.000 |
| CM-Diverse | 0.773 | 0.056 | 0.008 | -1.722[c] | 30.650 | 0.095 | 0.381 |
| **VPL and other thalamic nuclei** | | | | | | | |
| VPL-Closed loop | 0.037 | 0.192 | 0.657 | 464.000[b] | - | 0.147 | 0.589 |
| VPL-Forked | 0.006 | 0.428 | 0.934 | 366.000[b] | - | 0.239 | 0.957 |
| VPL-L-shape | 0.246 | 0.677 | 0.165 | -0.758[a] | 38.000 | 0.453 | 1.000 |
| VPL-Diverse | 0.249 | 0.480 | 0.094 | -1.547[a] | 38.000 | 0.130 | 0.521 |

a = t statistic from Student's t-test; b = W statistic from Mann-Whitney U test; c = t statistic from Welch's t-test; asterisks indicate significance after Bonferroni correction; all tests were two-sided.

**Table S20. Effect of sleep stages on fractions of thalamo-cortical motif types in Sleep Dataset-WU**

| Motif | F | DF1 | DF2 | p | $p_{Bonf}$ |
|---|---|---|---|---|---|
| **CM and other cortical regions** | | | | | |
| CM-Closed loop | 7.871 | 2.000 | 36.387 | 0.001 | 0.006** |
| CM-Forked | 0.020 | 2.000 | 33.176 | 0.981 | 1.000 |
| CM-L-shape | 0.983 | 2.000 | 29.608 | 0.386 | 1.000 |
| CM-Diverse | 1.724 | 2.000 | 40.947 | 0.191 | 0.764 |
| **VPL and other cortical regions** | | | | | |
| VPL-Closed loop | 5.488 | 2.000 | 32.546 | 0.009 | 0.035* |
| VPL-Forked | 1.634 | 2.000 | 35.108 | 0.210 | 0.838 |
| VPL-L-shape | 1.200 | 2.000 | 33.035 | 0.314 | 1.000 |
| VPL-Diverse | 1.441 | 2.000 | 38.659 | 0.249 | 0.997 |

Asterisks indicate significance after Bonferroni correction.

**Table S21. Post hoc comparisons for thalamo-cortical motif types with significant sleep-stage effects in Sleep Dataset-WU**

| Motif | Level 1 | Level 2 | Estimate | SE | DF | z | p | $p_{Bonf}$ |
|---|---|---|---|---|---|---|---|---|
| **CM and other cortical regions** | | | | | | | | |
| CM-Closed loop | W | N2 | 0.029 | 0.009 | Inf | 3.093 | 0.002 | 0.006** |
| CM-Closed loop | W | N3 | 0.045 | 0.012 | Inf | 3.596 | < 0.001 | 0.001** |
| CM-Closed loop | N2 | N3 | 0.016 | 0.012 | Inf | 1.321 | 0.186 | 0.559 |
| **VPL and other cortical regions** | | | | | | | | |
| VPL-Closed loop | W | N2 | 0.020 | 0.008 | Inf | 2.432 | 0.015 | 0.045* |
| VPL-Closed loop | W | N3 | 0.035 | 0.011 | Inf | 3.093 | 0.002 | 0.006** |
| VPL-Closed loop | N2 | N3 | 0.015 | 0.011 | Inf | 1.381 | 0.167 | 0.501 |

Asterisks indicate significance after Bonferroni correction; all tests were two-sided.

**Table S22. Comparisons of fractions of thalamo-cortical motif types between wakefulness and deep sedation in Anesthesia Dataset-SHH**

| Motif | p for Shapiro-Wilk normality test | statistic | df | p | $p_{Bonf}$ |
|---|---|---|---|---|---|
| **CM and other cortical regions** | | | | | |
| CM-Closed loop | 0.008 | 178.000[b] | - | 0.435 | 1.000 |
| CM-Forked | 0.653 | 0.216[a] | 23.000 | 0.831 | 1.000 |
| CM-L-shape | 0.050 | 169.000[b] | - | 0.603 | 1.000 |
| CM-Diverse | 0.176 | -1.672[a] | 23.000 | 0.108 | 0.433 |
| **VPL and other cortical regions** | | | | | |
| VPL-Closed loop | 0.392 | 0.883[a] | 23.000 | 0.387 | 1.000 |
| VPL-Forked | 0.596 | -0.871[a] | 23.000 | 0.393 | 1.000 |
| VPL-L-shape | 0.292 | 2.860[a] | 23.000 | 0.009 | 0.035* |
| VPL-Diverse | 0.025 | 91.000[b] | - | 0.095 | 0.380 |

a = t statistic from Student's t-test; b = W statistic from Wilcoxon signed-rank test; asterisks indicate significance after Bonferroni correction; all tests were two-sided.

**Table S23. Comparisons of fractions of thalamo-cortical motif types between HC and UWS in UWS Dataset-ZJH**

| Motif | p for Shapiro-Wilk normality test | | p for Levene's test for equal variances | statistic | df | p | $p_{Bonf}$ |
|---|---|---|---|---|---|---|---|
| | HC | UWS | | | | | |
| **CM and other cortical regions** | | | | | | | |
| CM-Closed loop | 0.372 | 0.537 | 0.447 | 1.845[a] | 38.000 | 0.073 | 0.292 |
| CM-Forked | 0.026 | 0.001 | 0.156 | 364.000[b] | - | 0.221 | 0.884 |
| CM-L-shape | 0.370 | 0.589 | 0.193 | 1.645[a] | 38.000 | 0.108 | 0.433 |

| | | | | | | | |
|---|---|---|---|---|---|---|---|
| CM-Diverse | 0.669 | 0.001 | 0.104 | 387.000[b] | - | 0.547 | 1.000 |
| **VPL and other cortical regions** | | | | | | | |
| VPL-Closed loop | 0.496 | 0.086 | 0.359 | 1.574[a] | 38.000 | 0.124 | 0.496 |
| VPL-Forked | 0.021 | 0.070 | 0.867 | 364.000[b] | - | 0.221 | 0.884 |
| VPL-L-shape | 0.869 | 0.652 | 0.339 | 1.261[a] | 38.000 | 0.215 | 0.861 |
| VPL-Diverse | 0.085 | 0.026 | 0.468 | 390.000[b] | - | 0.602 | 1.000 |

a = t statistic from Student's t-test; b = W statistic from Mann–Whitney U test; asterisks indicate significance after Bonferroni correction; all tests were two-sided.

**Table S24. Effect of sleep stages on effective connectivity along thalamic pathways in Sleep Dataset-WU**

| Pathway | F | DF1 | DF2 | p | $p_{Bonf}$ |
|---|---|---|---|---|---|
| CMtoVPL | 5.817 | 2.000 | 34.715 | 0.007 | 0.013* |
| VPLtoPUL | 7.132 | 2.000 | 34.735 | 0.003 | 0.005** |

Asterisks indicate significance after Bonferroni correction.

**Table S25. Post hoc comparisons for thalamic pathways showing significant sleep-stage effects in Sleep Dataset-WU**

| Pathway | Level 1 | Level 2 | Estimate | SE | DF | z | p | $p_{Bonf}$ |
|---|---|---|---|---|---|---|---|---|
| CMtoVPL | W | N2 | -0.023 | 0.017 | Inf | -1.404 | 0.160 | 0.481 |
| CMtoVPL | W | N3 | -0.074 | 0.022 | Inf | -3.409 | 0.001 | 0.002** |
| CMtoVPL | N2 | N3 | -0.051 | 0.021 | Inf | -2.469 | 0.014 | 0.041* |
| VPLtoPUL | W | N2 | 0.003 | 0.015 | Inf | 0.182 | 0.856 | 1.000 |
| VPLtoPUL | W | N3 | -0.066 | 0.020 | Inf | -3.304 | 0.001 | 0.003** |
| VPLtoPUL | N2 | N3 | -0.068 | 0.019 | Inf | -3.628 | < 0.001 | 0.001** |

Asterisks indicate significance after Bonferroni correction; all tests were two-sided.

**Table S26. Comparisons of effective connectivity along thalamic pathways between wakefulness and deep sedation in Anesthesia Dataset-SHH**

| Pathway | p for Shapiro–Wilk normality test | statistic | df | p | $p_{Bonf}$ |
|---|---|---|---|---|---|
| CMtoVPL | 0.381 | -0.774[a] | 23 | 0.447 | 0.893 |
| VPLtoPUL | 0.977 | -1.879[a] | 23 | 0.073 | 0.146 |

a = t statistic from Student's t-test; asterisks indicate significance after Bonferroni correction; all tests were two-sided.

**Table S27. Comparisons of effective connectivity along thalamic pathways between HC and UWS in UWS Dataset-ZJH**

| Pathway | p for Shapiro-Wilk | p for Levene's test for | statistic | df | p | $p_{Bonf}$ |
|---|---|---|---|---|---|---|

| | normality test | | equal variances | | | | |
|---|---|---|---|---|---|---|---|
| | HC | UWS | | | | | |
| CMtoVPL | 0.072 | 0.239 | 0.080 | 0.620[a] | 38 | 0.539 | 1.000 |
| VPLtoPUL | 0.230 | 0.219 | 0.775 | -0.150[a] | 38 | 0.881 | 1.000 |

a = t statistic from Student's t-test; asterisks indicate significance after Bonferroni correction; all tests were two-sided.

**Table S28. Effect of sleep stages on thalamic community similarity in Sleep Dataset-IUGM**

| ROI | F | DF1 | DF2 | p | $p_{Bonf}$ |
|---|---|---|---|---|---|
| CM | 10.060 | 2.000 | 17.923 | 0.001 | 0.002** |
| VPL | 7.666 | 2.000 | 17.416 | 0.004 | 0.008** |

Asterisks indicate significance after Bonferroni correction.

**Table S29. Post hoc comparisons for thalamic community similarity with significant sleep-stage effects in Sleep Dataset-IUGM**

| ROI | Level 1 | Level 2 | Estimate | SE | DF | z | p | $p_{Bonf}$ |
|---|---|---|---|---|---|---|---|---|
| CM | W | N2 | -0.273 | 0.071 | Inf | -3.845 | < 0.001 | < 0.001*** |
| CM | W | N3 | -0.298 | 0.079 | Inf | -3.782 | < 0.001 | < 0.001*** |
| CM | N2 | N3 | -0.026 | 0.079 | Inf | -0.323 | 0.746 | 1.000 |
| VPL | W | N2 | -0.176 | 0.066 | Inf | -2.658 | 0.008 | 0.024* |
| VPL | W | N3 | -0.277 | 0.074 | Inf | -3.763 | < 0.001 | 0.001** |
| VPL | N2 | N3 | -0.101 | 0.074 | Inf | -1.376 | 0.169 | 0.507 |

Asterisks indicate significance after Bonferroni correction; all tests were two-sided.

**Table S30. Effect of sleep stages on the fraction of thalamic-thalamic motif types in Sleep Dataset-IUGM**

| Motif | F | DF1 | DF2 | p | $p_{Bonf}$ |
|---|---|---|---|---|---|
| **CM and other thalamic nuclei** | | | | | |
| CM-Closed loop | 3.767 | 2.000 | 17.019 | 0.044 | 0.177 |
| CM-Forked | 5.096 | 2.000 | 18.002 | 0.018 | 0.071 |
| CM-L-shape | 1.133 | 2.000 | 18.757 | 0.343 | 1.000 |
| CM-Diverse | 3.007 | 2.000 | 14.051 | 0.082 | 0.327 |
| **VPL and other thalamic nuclei** | | | | | |
| VPL-Closed loop | 2.398 | 2.000 | 18.071 | 0.119 | 0.477 |
| VPL-Forked | 0.247 | 2.000 | 18.164 | 0.784 | 1.000 |
| VPL-L-shape | 2.474 | 2.000 | 28.000 | 0.102 | 0.410 |
| VPL-Diverse | 2.675 | 2.000 | 16.111 | 0.099 | 0.397 |

Asterisks indicate significance after Bonferroni correction.

**Table S31. Post hoc comparisons for thalamic-thalamic motif types in Sleep Dataset-IUGM**

| Motif | Level 1 | Level 2 | Estimate | SE | DF | z | p | $p_{Bonf}$ |
|---|---|---|---|---|---|---|---|---|
| **CM and other thalamic nuclei** | | | | | | | | |
| CM-Closed loop | W | N2 | -0.189 | 0.113 | Inf | -1.678 | 0.093 | 0.280 |
| CM-Closed loop | W | N3 | -0.339 | 0.126 | Inf | -2.698 | 0.007 | 0.021* |
| CM-Closed loop | N2 | N3 | -0.150 | 0.125 | Inf | -1.199 | 0.230 | 0.691 |
| CM-Forked | W | N2 | 0.093 | 0.034 | Inf | 2.756 | 0.006 | 0.018* |
| CM-Forked | W | N3 | 0.101 | 0.038 | Inf | 2.698 | 0.007 | 0.021* |
| CM-Forked | N2 | N3 | 0.008 | 0.037 | Inf | 0.226 | 0.821 | 1.000 |
| CM-L-shape | W | N2 | 0.030 | 0.066 | Inf | 0.459 | 0.646 | 1.000 |
| CM-L-shape | W | N3 | 0.109 | 0.074 | Inf | 1.483 | 0.138 | 0.415 |
| CM-L-shape | N2 | N3 | 0.079 | 0.072 | Inf | 1.088 | 0.277 | 0.830 |
| CM-Diverse | W | N2 | 0.070 | 0.048 | Inf | 1.455 | 0.146 | 0.437 |
| CM-Diverse | W | N3 | 0.129 | 0.054 | Inf | 2.417 | 0.016 | 0.047* |
| CM-Diverse | N2 | N3 | 0.059 | 0.054 | Inf | 1.108 | 0.268 | 0.803 |
| **VPL and other thalamic nuclei** | | | | | | | | |
| VPL-Closed loop | W | N2 | -0.027 | 0.112 | Inf | -0.244 | 0.807 | 1.000 |
| VPL-Closed loop | W | N3 | -0.254 | 0.124 | Inf | -2.041 | 0.041 | 0.124 |
| VPL-Closed loop | N2 | N3 | -0.226 | 0.122 | Inf | -1.847 | 0.065 | 0.194 |
| VPL-Forked | W | N2 | 0.017 | 0.028 | Inf | 0.614 | 0.539 | 1.000 |
| VPL-Forked | W | N3 | 0.018 | 0.031 | Inf | 0.586 | 0.558 | 1.000 |
| VPL-Forked | N2 | N3 | 0.001 | 0.031 | Inf | 0.035 | 0.972 | 1.000 |
| VPL-L-shape | W | N2 | -0.018 | 0.062 | Inf | -0.292 | 0.770 | 1.000 |
| VPL-L-shape | W | N3 | 0.125 | 0.069 | Inf | 1.811 | 0.070 | 0.210 |
| VPL-L-shape | N2 | N3 | 0.143 | 0.068 | Inf | 2.111 | 0.035 | 0.104 |
| VPL-Diverse | W | N2 | 0.030 | 0.043 | Inf | 0.679 | 0.497 | 1.000 |
| VPL-Diverse | W | N3 | 0.110 | 0.048 | Inf | 2.275 | 0.023 | 0.069 |
| VPL-Diverse | N2 | N3 | 0.081 | 0.048 | Inf | 1.684 | 0.092 | 0.277 |

Asterisks indicate significance after Bonferroni correction; all tests were two-sided.

**Table S32. Effect of sleep stages on effective connectivity along thalamic pathways in Sleep Dataset-IUGM**

| Pathway | F | DF1 | DF2 | p | $p_{Bonf}$ |
|---|---|---|---|---|---|
| CMtoVPL | 1.474 | 2.000 | 21.037 | 0.252 | 0.504 |
| VPLtoPUL | 2.265 | 2.000 | 20.464 | 0.129 | 0.259 |

Asterisks indicate significance after Bonferroni correction.

**Table S33. Post hoc comparisons for thalamic pathways in Sleep Dataset-IUGM**

| Pathway | Level 1 | Level 2 | Estimate | SE | DF | z | p | $p_{Bonf}$ |
|---|---|---|---|---|---|---|---|---|
| CMtoVPL | W | N2 | 0.030 | 0.029 | Inf | 1.041 | 0.298 | 0.893 |
| CMtoVPL | W | N3 | -0.023 | 0.032 | Inf | -0.727 | 0.467 | 1.000 |

| | | | | | | | | |
|---|---|---|---|---|---|---|---|---|
| CMtoVPL | N2 | N3 | -0.053 | 0.031 | Inf | -1.684 | 0.092 | 0.277 |
| VPLtoPUL | W | N2 | 0.006 | 0.025 | Inf | 0.223 | 0.824 | 1.000 |
| VPLtoPUL | W | N3 | -0.049 | 0.028 | Inf | -1.777 | 0.075 | 0.226 |
| VPLtoPUL | N2 | N3 | -0.054 | 0.027 | Inf | -1.997 | 0.046 | 0.138 |

Asterisks indicate significance after Bonferroni correction; all tests were two-sided.

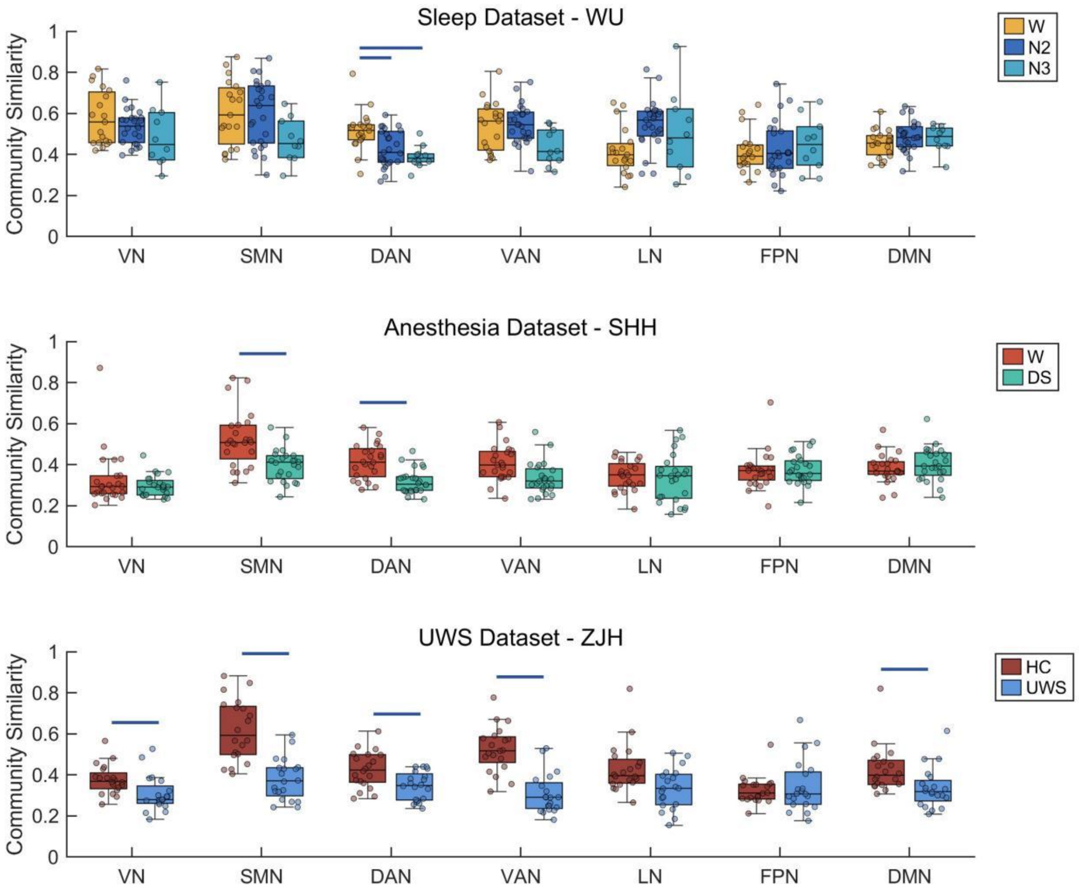


Figure S1. Comparison of network-wise community similarity across conditions in Sleep Dataset-WU, Anesthesia Dataset-SHH, and UWS Dataset-ZJH. Blue line denotes significant decreases at $p < 0.05$ (Bonferroni-corrected). Center line indicates median, box indicates interquartile range, and whisker denotes 1.5 × IQR. W = wakefulness; DS = deep sedation; HC = healthy controls; UWS = unresponsive wakefulness syndrome.

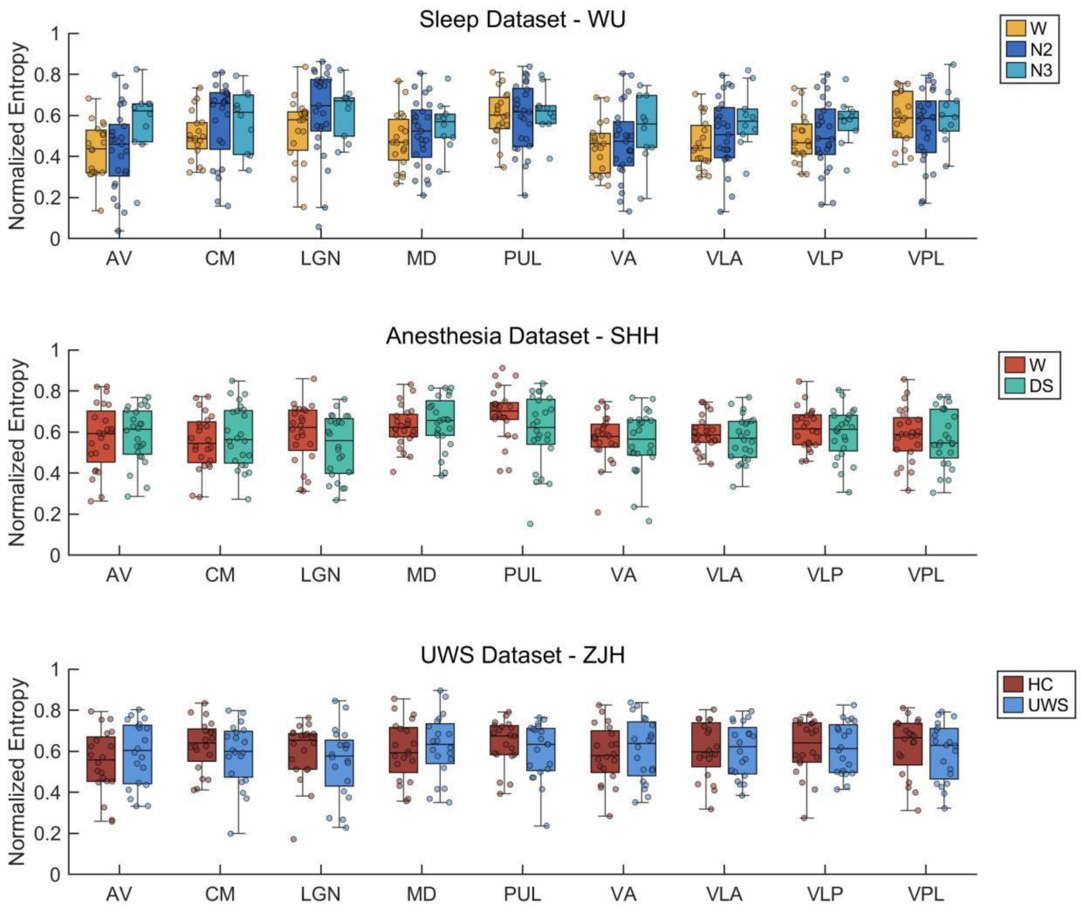


Figure S2. Comparison of thalamic normalized entropy across conditions in Sleep Dataset-WU, Anesthesia Dataset-SHH, and UWS Dataset-ZJH. Center line indicates median, box indicates interquartile range, and whisker denotes 1.5 × IQR. W = wakefulness; DS = deep sedation; HC = healthy controls; UWS = unresponsive wakefulness syndrome.

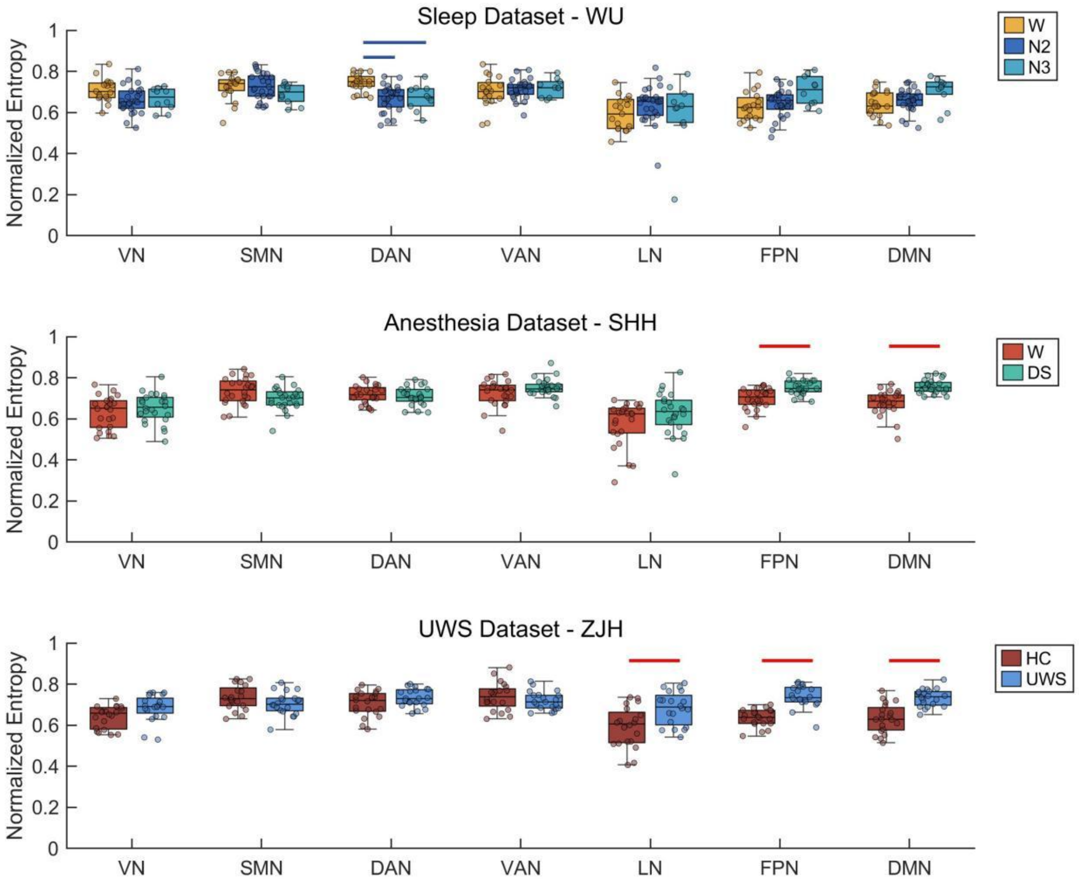


Figure S3. Comparison of network-wise normalized entropy across conditions in Sleep Dataset-WU, Anesthesia Dataset-SHH, and UWS Dataset-ZJH. Red line denotes significant increases and blue line denotes significant decreases at $p < 0.05$ (Bonferroni-corrected). Center line indicates median, box indicates interquartile range, and whisker denotes 1.5 × IQR. W = wakefulness; DS = deep sedation; HC = healthy controls; UWS = unresponsive wakefulness syndrome.

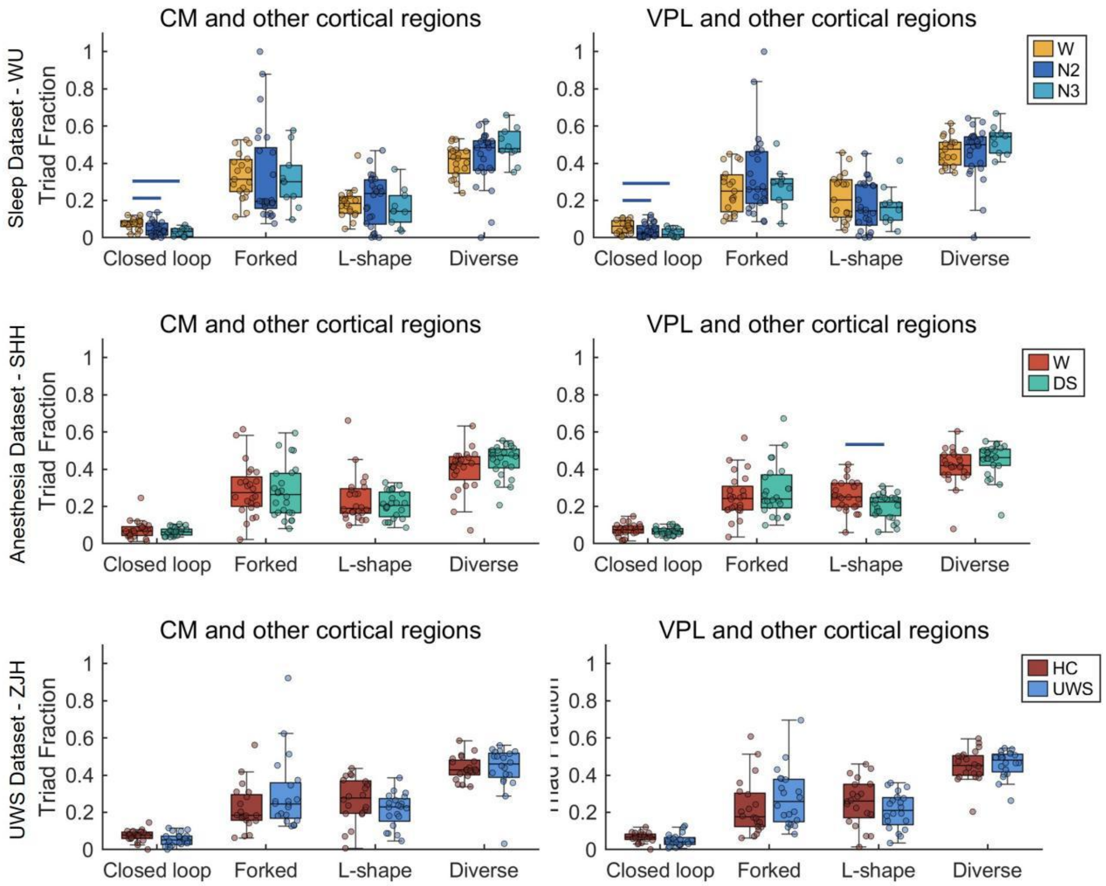


Figure S4. Comparison of the fraction of thalamic-cortical motif types centered on CM and VPL across conditions in Sleep Dataset-WU, Anesthesia Dataset-SHH, and UWS Dataset-ZJH. Rows correspond to the datasets. Red line denotes significant increases and blue line denotes significant decreases at $p < 0.05$ (Bonferroni-corrected). Center line indicates median, box indicates interquartile range, and whisker denotes 1.5 × IQR. W = wakefulness; DS = deep sedation; HC = healthy controls; UWS = unresponsive wakefulness syndrome.

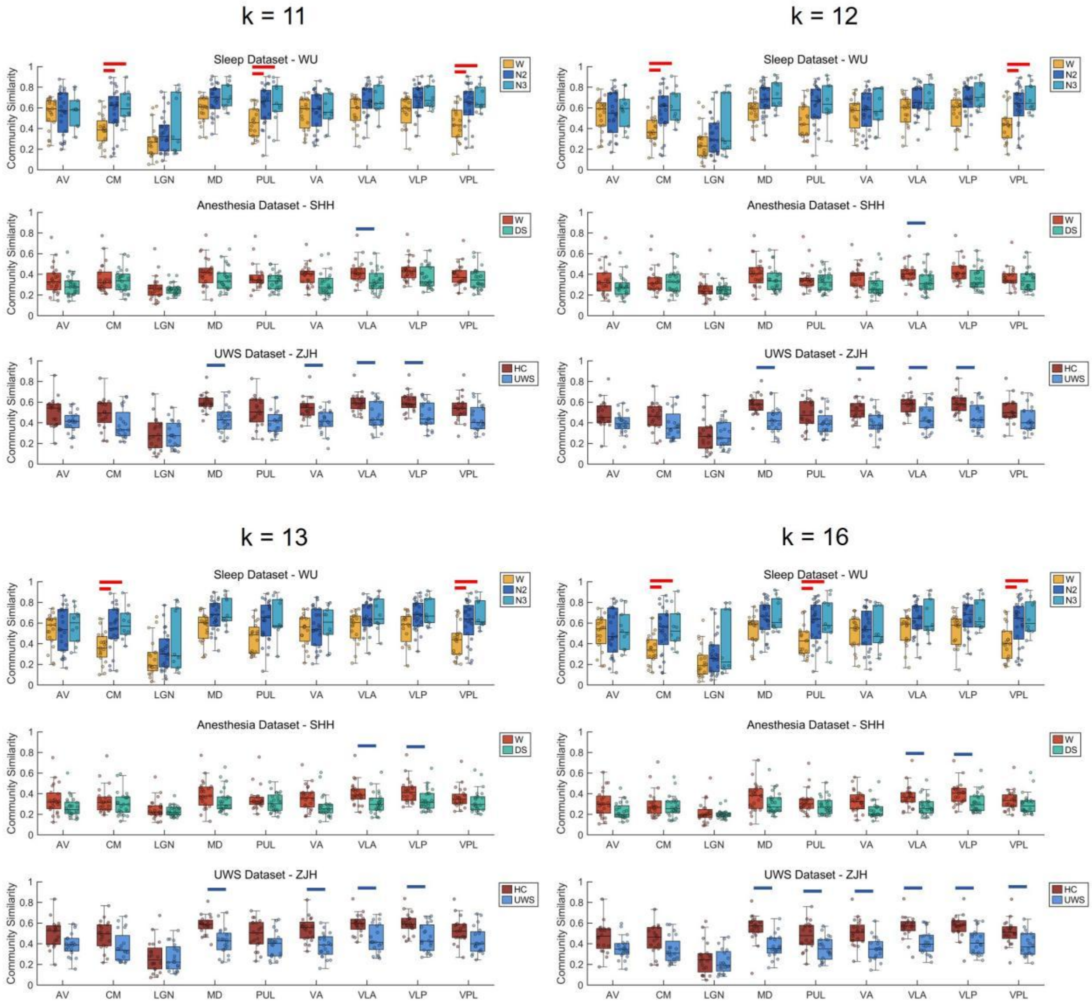


Figure S5. Comparison of thalamic community similarity across conditions in Sleep Dataset-WU, Anesthesia Dataset-SHH, and UWS Dataset-ZJH with k = 11, 12, 13, 16. Red line denotes significant increases and blue line denotes significant decreases at $p < 0.05$ (Bonferroni-corrected). Center line indicates median, box indicates interquartile range, and whisker denotes 1.5 × IQR. W = wakefulness; DS = deep sedation; HC = healthy controls; UWS = unresponsive wakefulness syndrome.

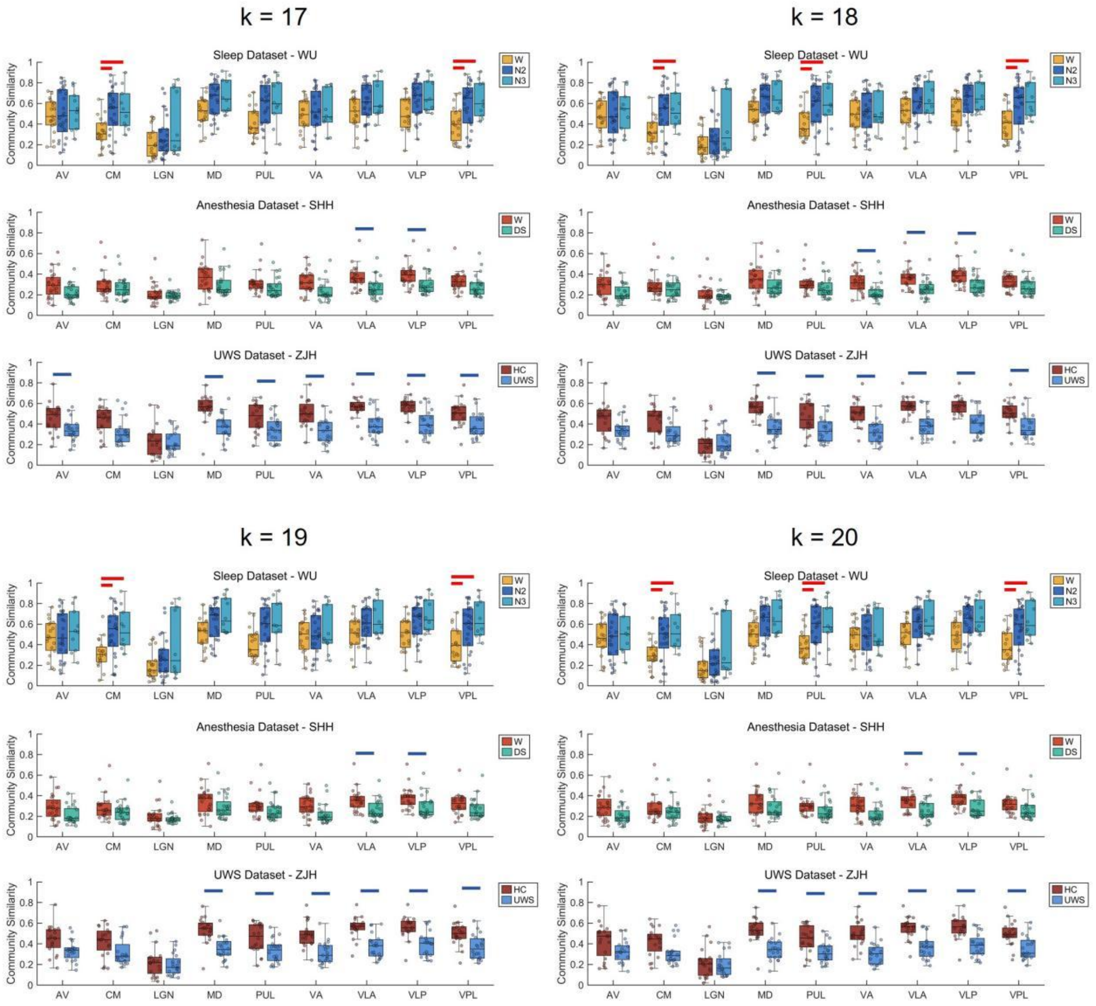


Figure S6. Comparison of thalamic community similarity across conditions in Sleep Dataset-WU, Anesthesia Dataset-SHH, and UWS Dataset-ZJH with k = 17, 18, 19, 20. Red line denotes significant increases and blue line denotes significant decreases at $p < 0.05$ (Bonferroni-corrected). Center line indicates median, box indicates interquartile range, and whisker denotes 1.5 × IQR. W = wakefulness; DS = deep sedation; HC = healthy controls; UWS = unresponsive wakefulness syndrome.

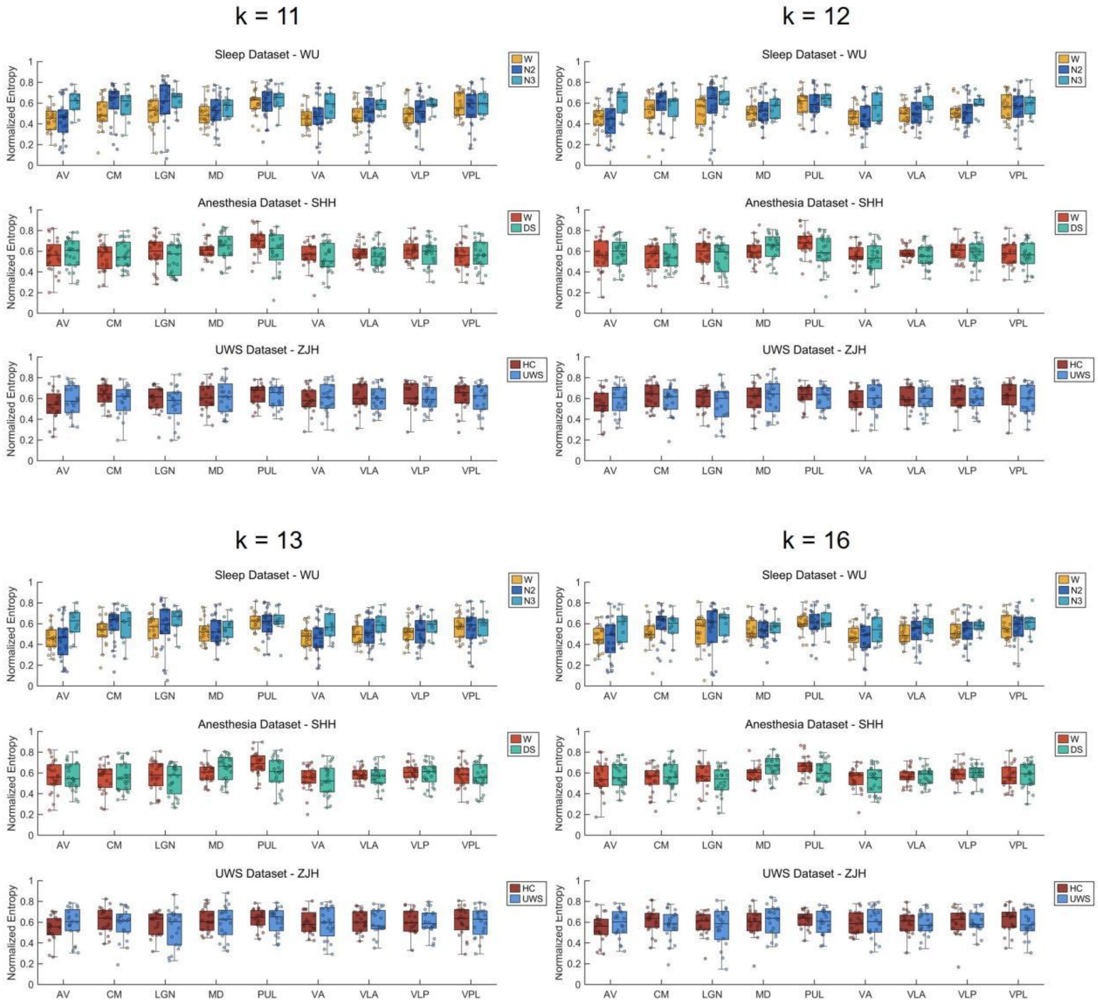


Figure S7. Comparison of thalamic normalized entropy across conditions in Sleep Dataset-WU, Anesthesia Dataset-SHH, and UWS Dataset-ZJH with k = 11, 12, 13, 16. Center line indicates median, box indicates interquartile range, and whisker denotes 1.5 × IQR. W = wakefulness; DS = deep sedation; HC = healthy controls; UWS = unresponsive wakefulness syndrome.

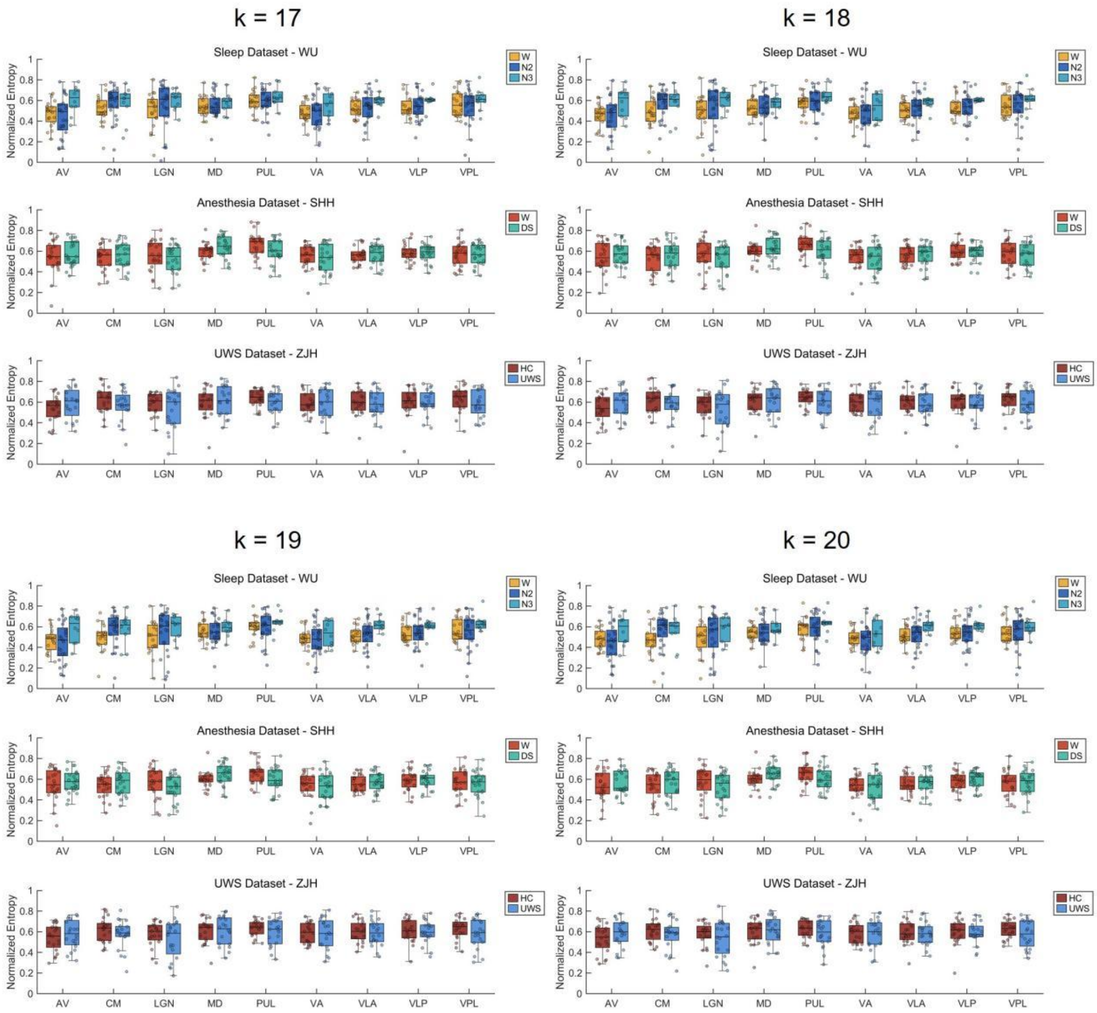


Figure S8. Comparison of thalamic normalized entropy across conditions in Sleep Dataset-WU, Anesthesia Dataset-SHH, and UWS Dataset-ZJH with k = 17, 18, 19, 20. Center line indicates median, box indicates interquartile range, and whisker denotes 1.5 × IQR. W = wakefulness; DS = deep sedation; HC = healthy controls; UWS = unresponsive wakefulness syndrome.

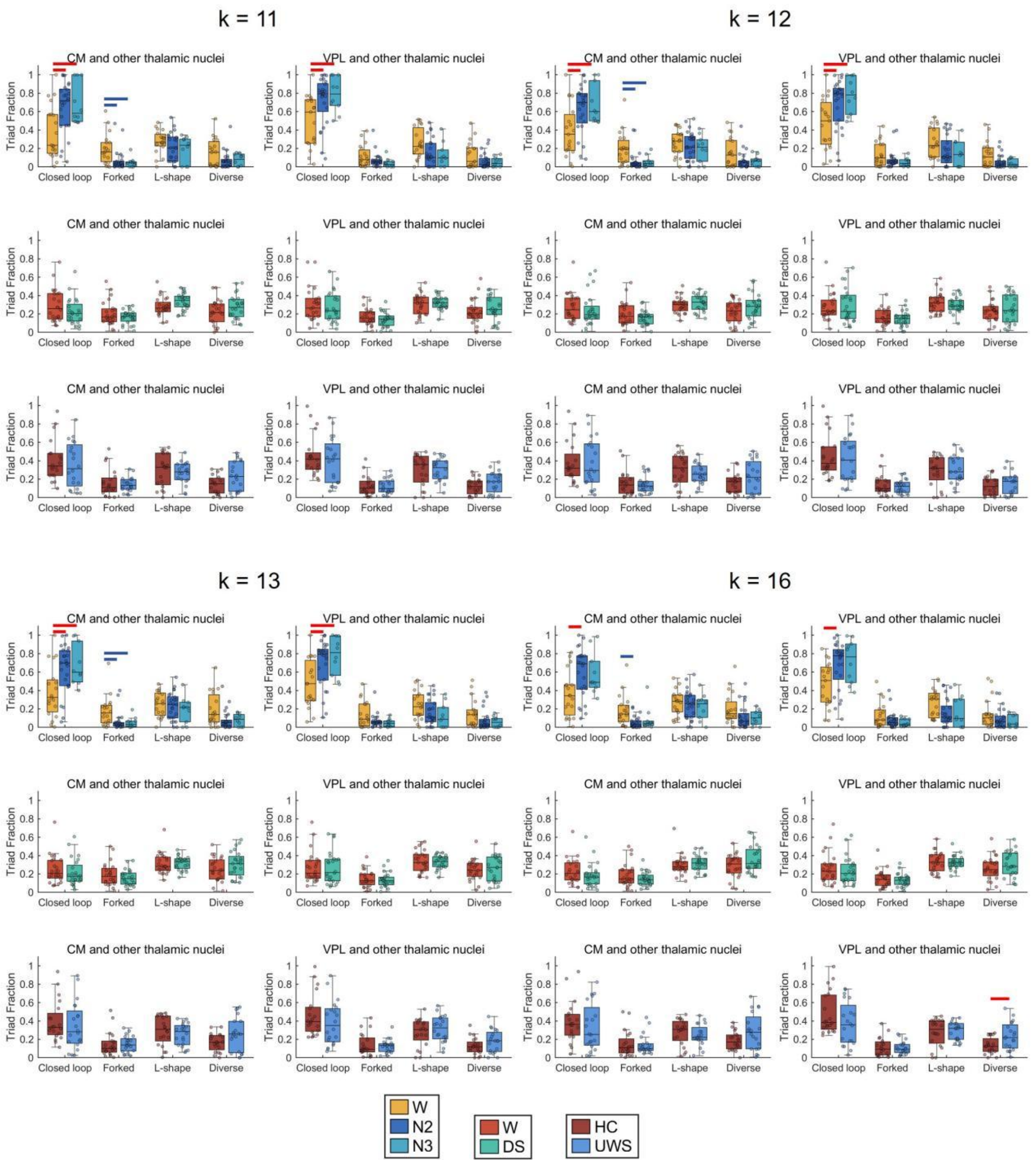

Figure S9. Comparison of the fraction of thalamic-thalamic motif types centered on CM and VPL across conditions in Sleep Dataset-WU, Anesthesia Dataset-SHH, and UWS Dataset-ZJH with k = 11, 12, 13, 16. Rows correspond to the datasets. Red line denotes significant increases and blue line denotes significant decreases at $p < 0.05$ (Bonferroni-corrected). Center line indicates median, box indicates interquartile range, and whisker denotes 1.5 × IQR. W = wakefulness; DS = deep sedation; HC = healthy controls; UWS = unresponsive wakefulness syndrome.

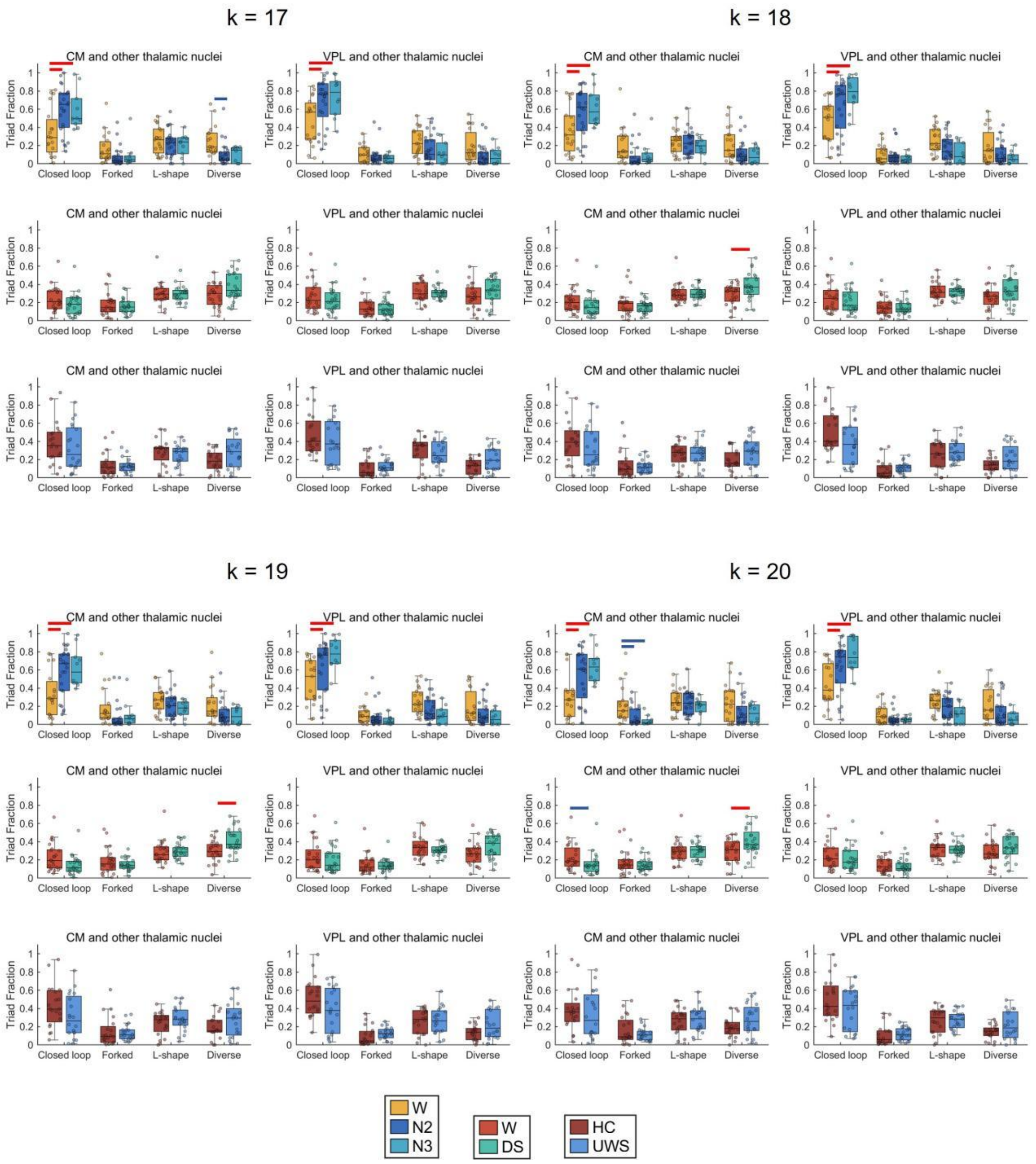


Figure S10. Comparison of the fraction of thalamic-thalamic motif types centered on CM and VPL across conditions in Sleep Dataset-WU, Anesthesia Dataset-SHH, and UWS Dataset-ZJH with k = 17, 18, 19, 20. Rows correspond to the datasets. Red line denotes significant increases and blue line denotes significant decreases at $p < 0.05$ (Bonferroni-corrected). Center line indicates median, box indicates interquartile range, and whisker denotes 1.5 × IQR. W = wakefulness; DS = deep sedation; HC = healthy controls; UWS = unresponsive wakefulness syndrome.

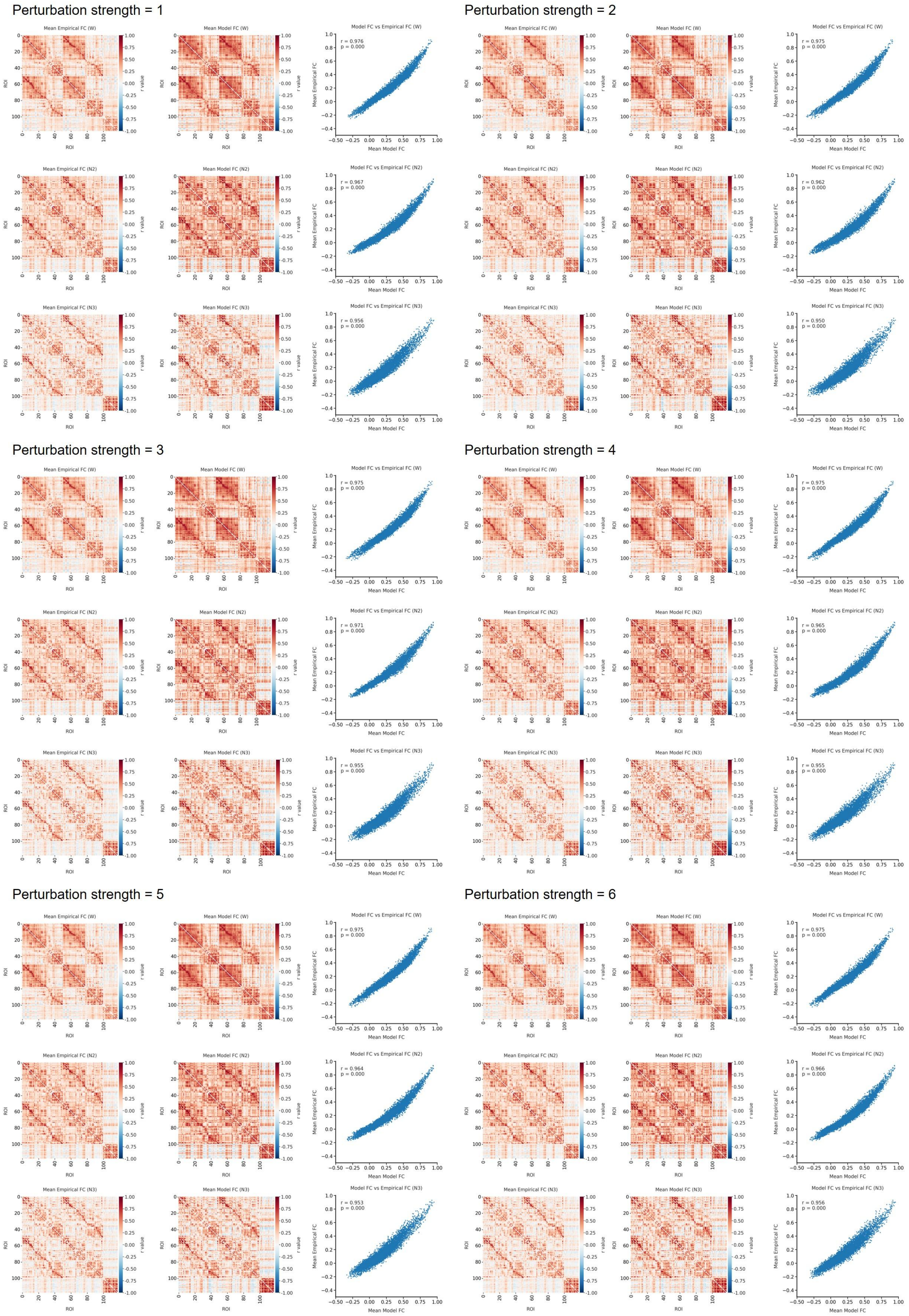


Figure S11. Similarity between mean empirical FC matrices and mean model-derived FC matrices across wakefulness (W), N2 sleep, and N3 sleep under perturbation strengths from 1 to 6 in Sleep Dataset-WU. Scatter plots show the relationship between empirical and model-derived FC values.

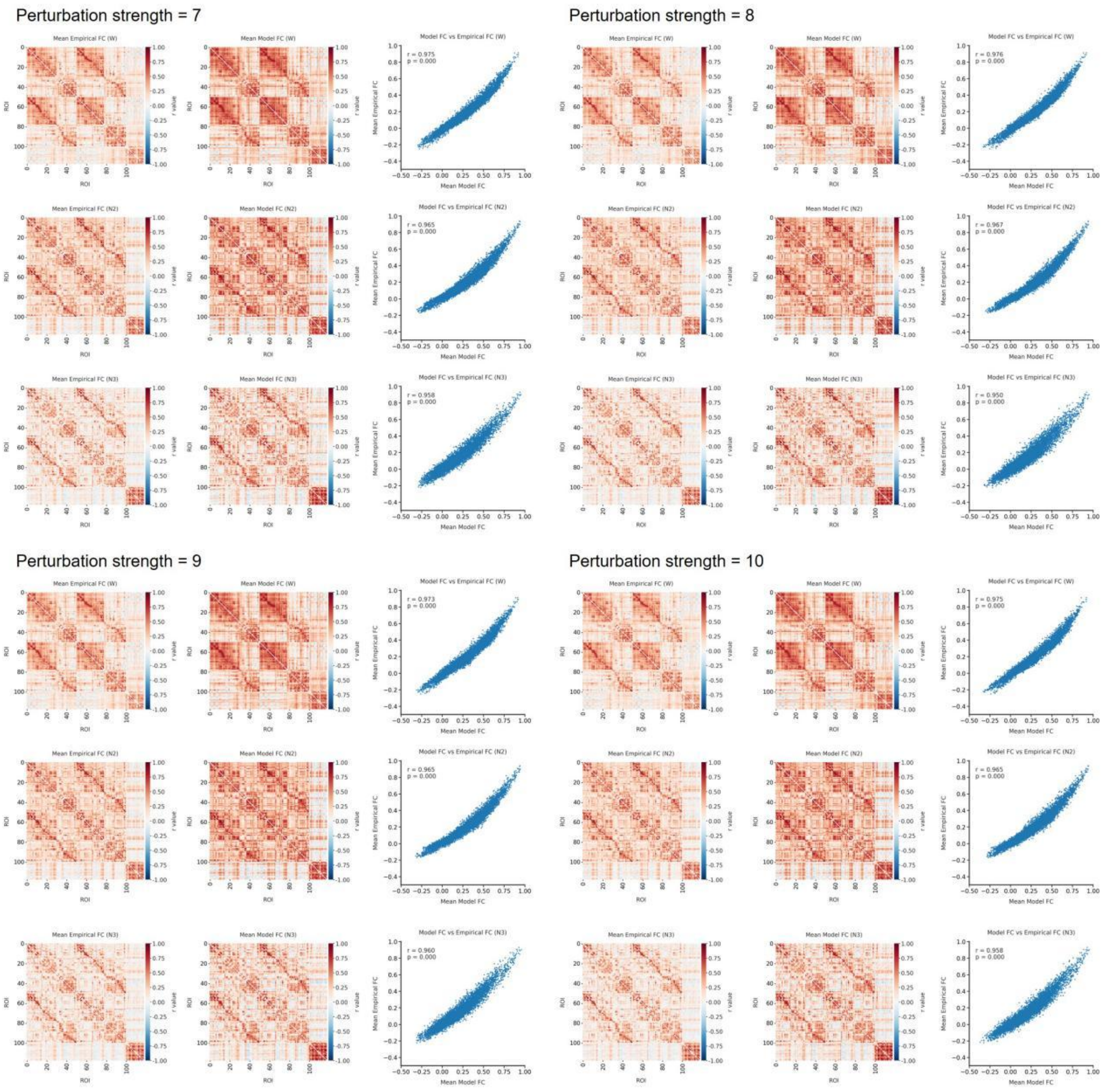


Figure S12. Similarity between mean empirical FC matrices and mean model-derived FC matrices across wakefulness (W), N2 sleep, and N3 sleep under perturbation strengths from 7 to 10 in Sleep Dataset-WU. Scatter plots show the relationship between empirical and model-derived FC values.

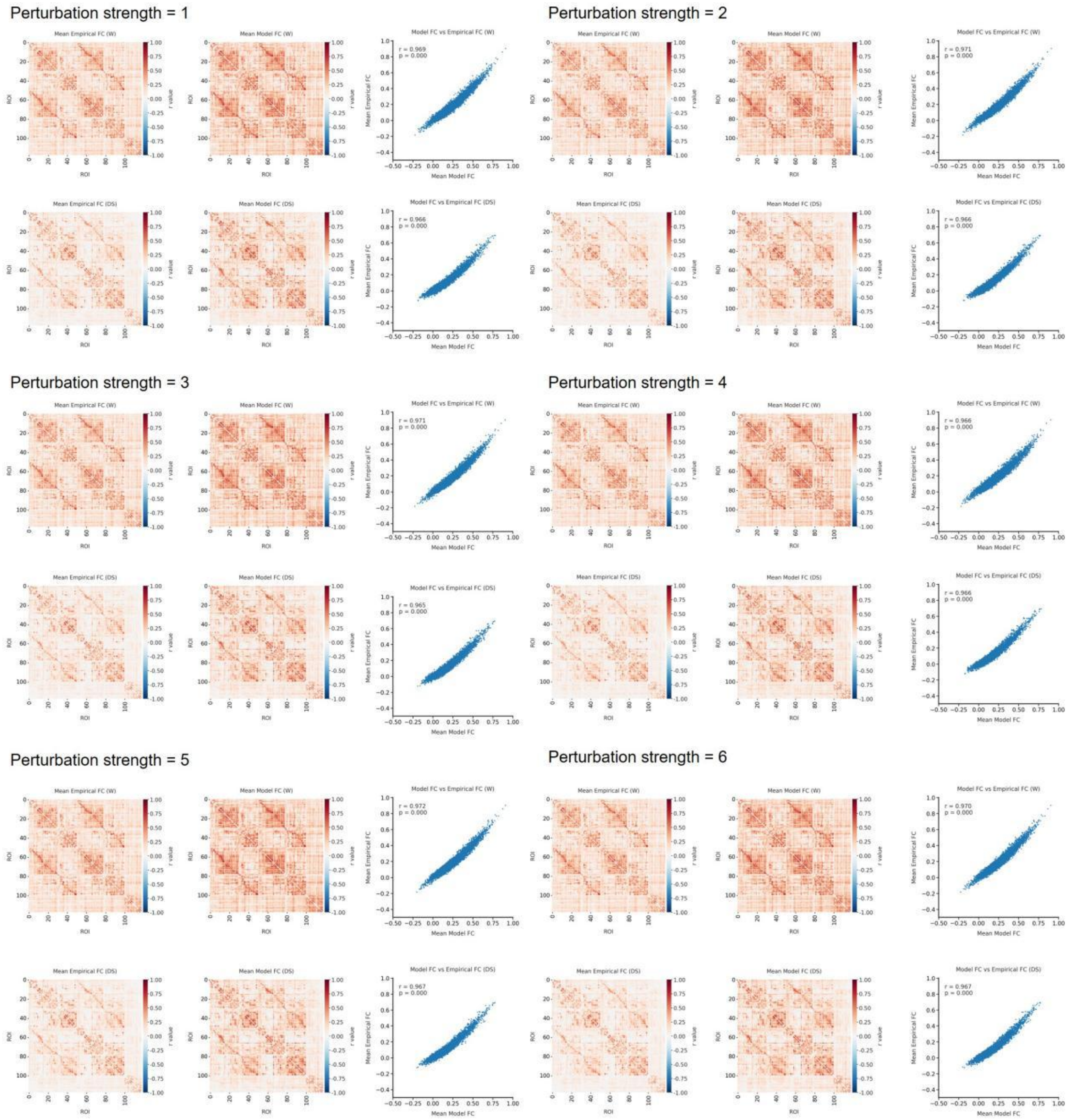


Figure S13. Similarity between mean empirical FC matrices and mean model-derived FC matrices across wakefulness (W) and deep sedation (DS) under perturbation strengths from 1 to 6 in Anesthesia Dataset-SHH. Scatter plots show the relationship between empirical and model-derived FC values.

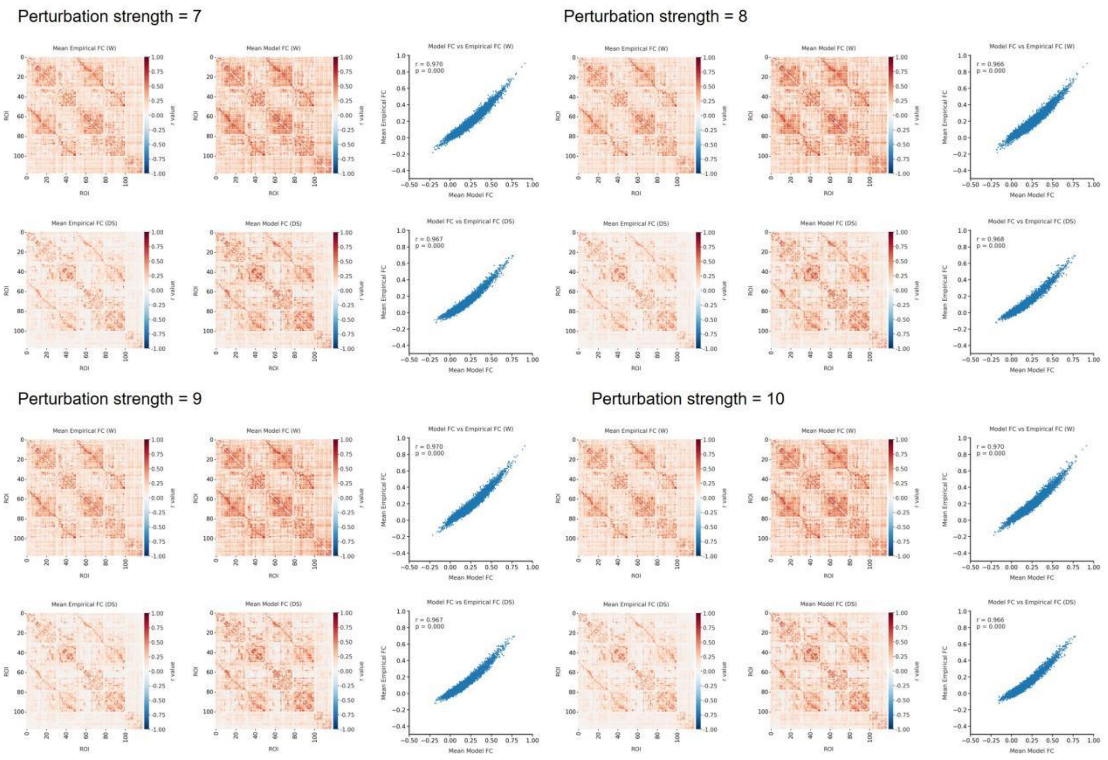


Figure S14. Similarity between mean empirical FC matrices and mean model-derived FC matrices across wakefulness (W) and deep sedation (DS) under perturbation strengths from 7 to 10 in Anesthesia Dataset-SHH. Scatter plots show the relationship between empirical and model-derived FC values.

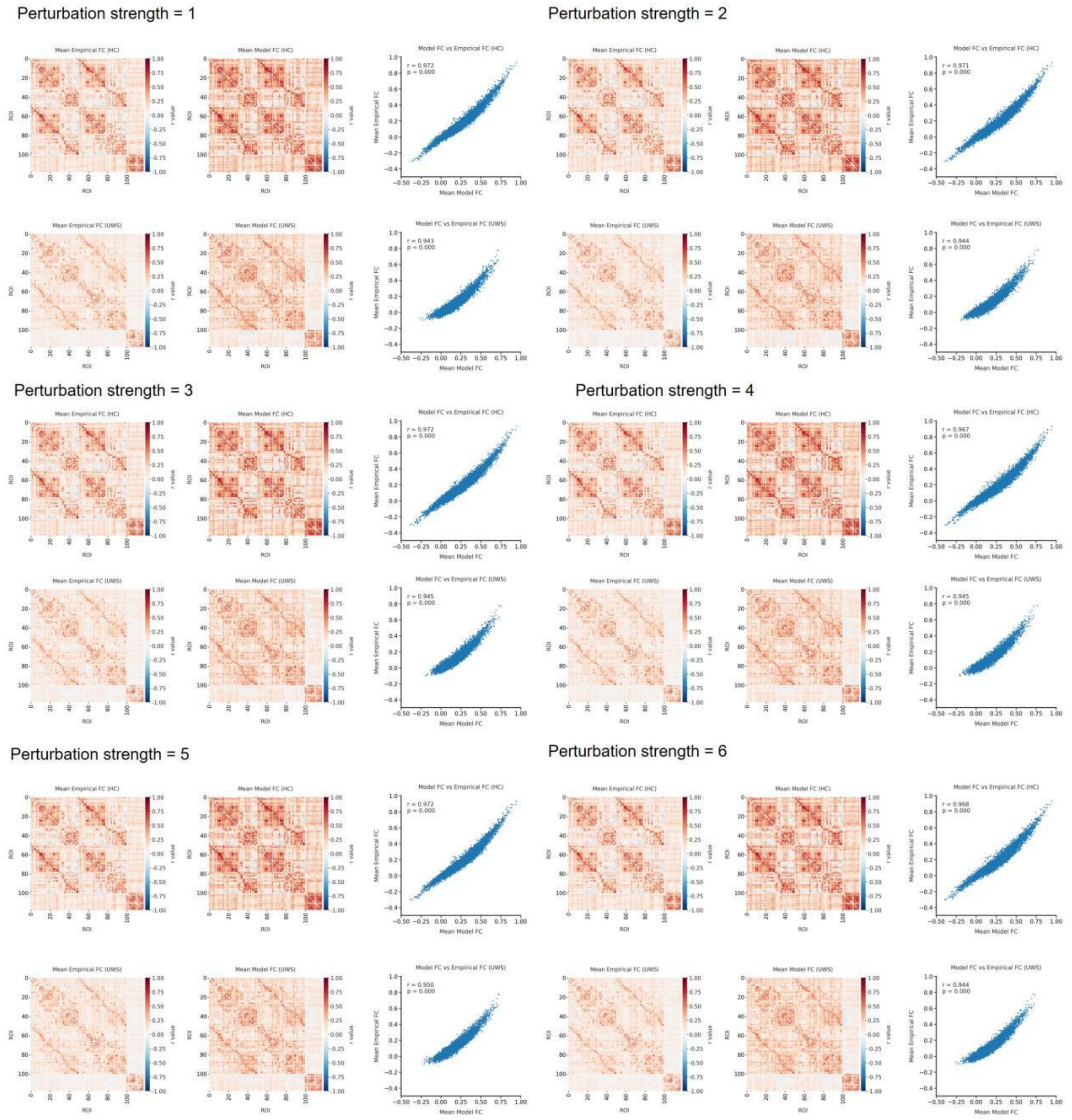


Figure S15. Similarity between mean empirical FC matrices and mean model-derived FC matrices across healthy controls (HC) and UWS under perturbation strengths from 1 to 6 in UWS Dataset-ZJH. Scatter plots show the relationship between empirical and model-derived FC values.

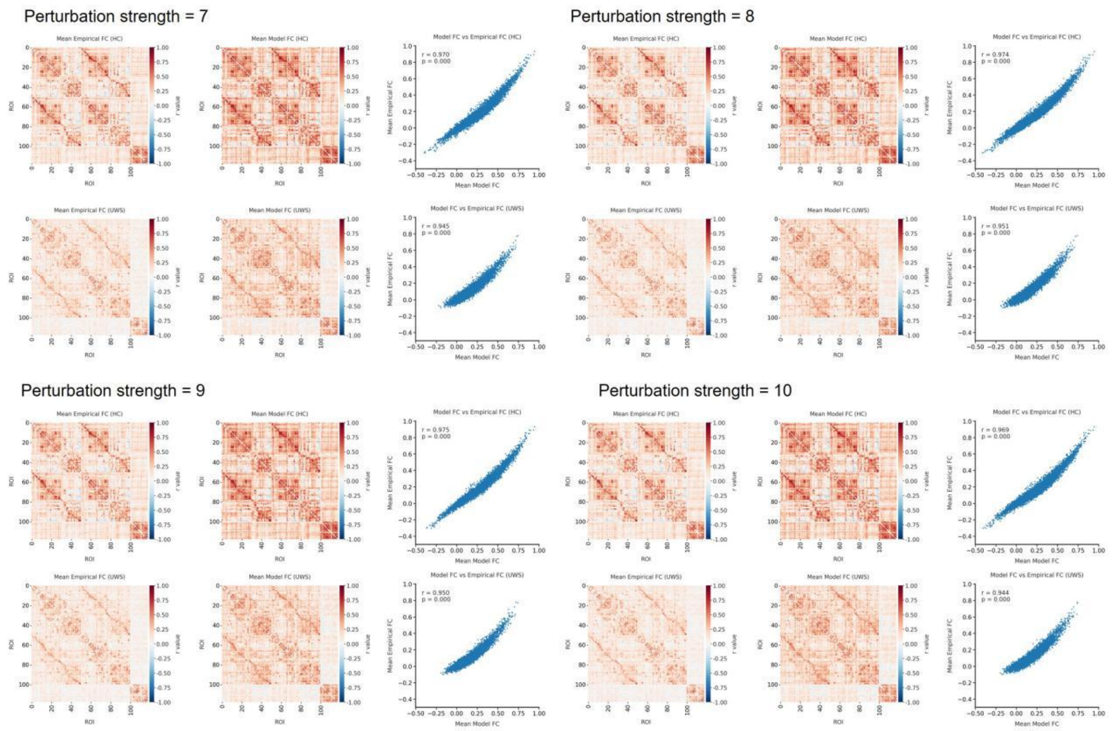


Figure S16. Similarity between mean empirical FC matrices and mean model-derived FC matrices across healthy controls (HC) and UWS under perturbation strengths from 7 to 10 in UWS Dataset-ZJH. Scatter plots show the relationship between empirical and model-derived FC values.

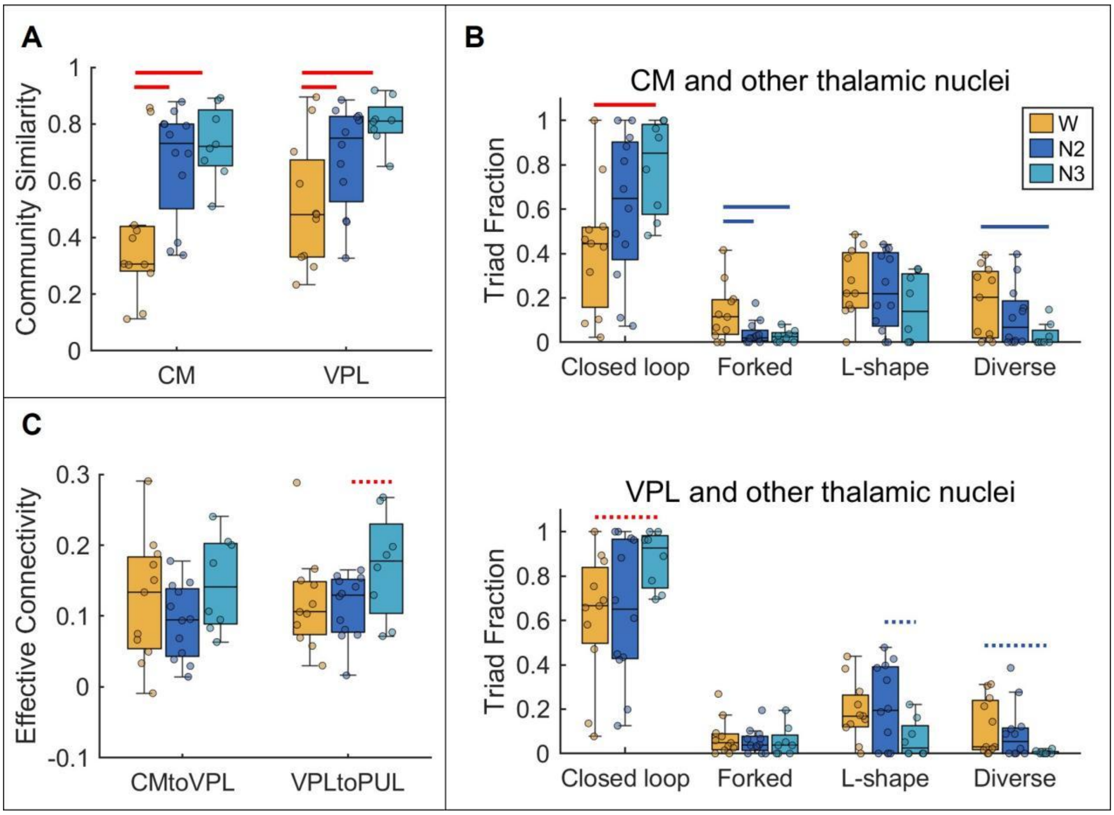


Figure S17. Comparison of sleep-specific thalamic coordination features across conditions in Sleep Dataset-IUGM. (A) Comparison of thalamic community similarity for CM and VPL. (B) Comparison of the fractions of thalamic-thalamic motif types for CM-centered and VPL-centered triads. (C) Comparison of effective connectivity for CM→VPL and VPL→PUL. Red solid line denotes significant increases and blue solid line denotes significant decreases at $p < 0.05$ (Bonferroni-corrected). Red dotted line indicate significant increase and blue dotted lines indicate significant decreases at $p < 0.05$ (uncorrected). Center line indicates median, box indicates interquartile range, and whisker denotes 1.5 × IQR. W = wakefulness.

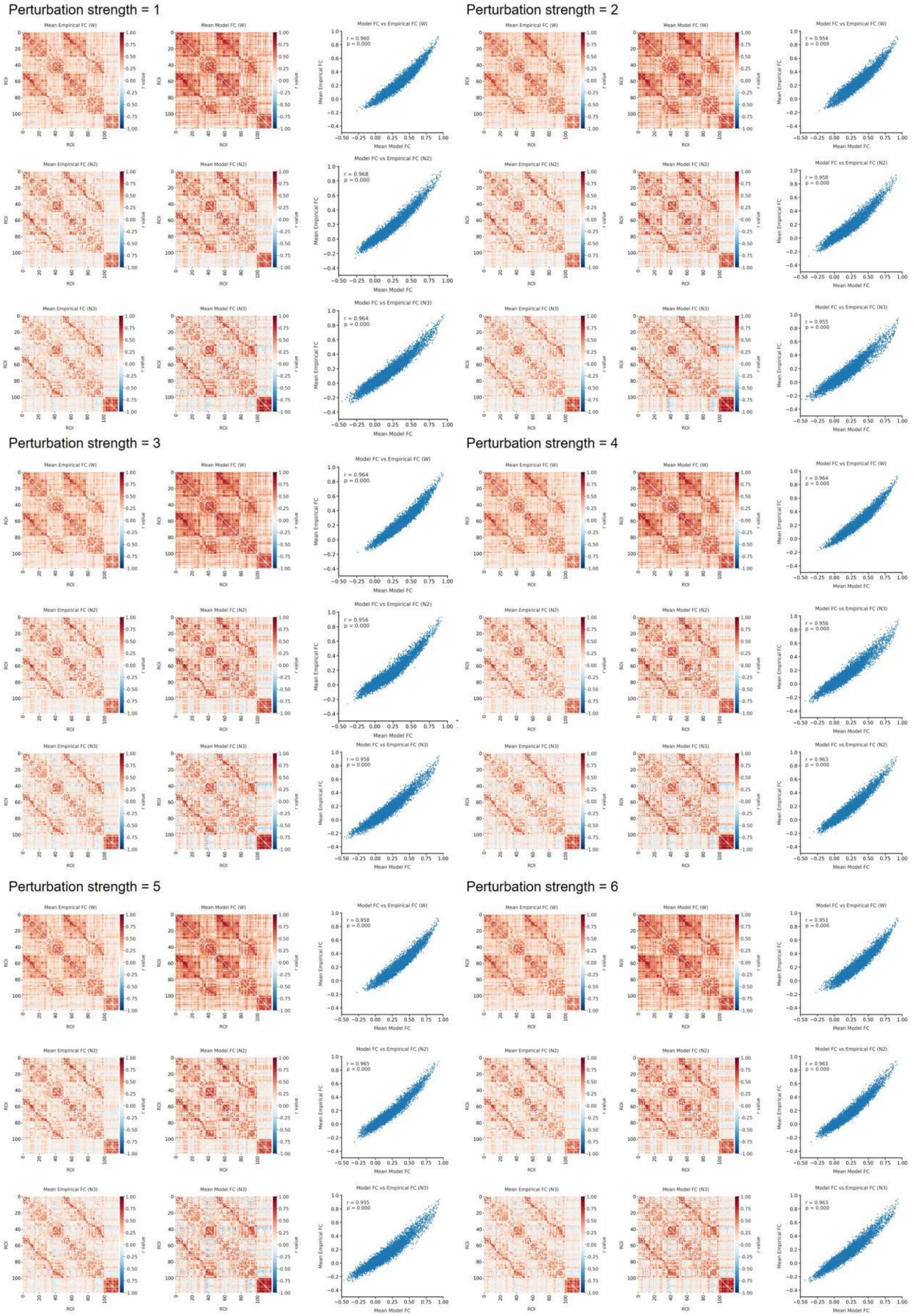


Figure S18. Similarity between mean empirical FC matrices and mean model-derived FC matrices across wakefulness (W), N2 sleep, and N3 sleep under perturbation strengths from 1 to 6 in Sleep Dataset-IUGM. Scatter plots show the relationship between empirical and model-derived FC values.

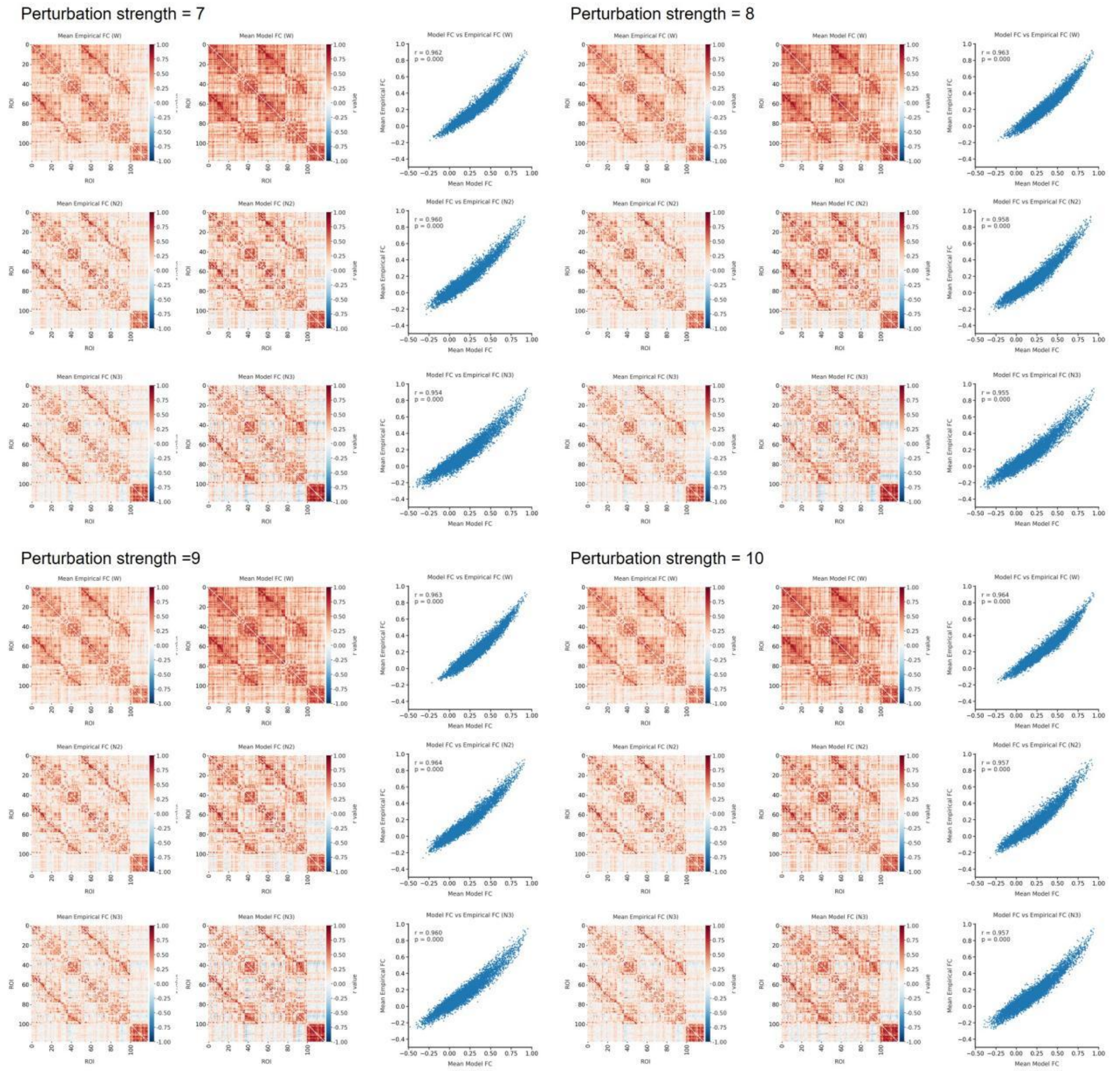


Figure S19. Similarity between mean empirical FC matrices and mean model-derived FC matrices across wakefulness (W), N2 sleep, and N3 sleep under perturbation strengths from 7 to 10 in Sleep Dataset-IUGM. Scatter plots show the relationship between empirical and model-derived FC values.
